# Design of a systemic small molecule clinical STING agonist using physics-based simulations and artificial intelligence

**DOI:** 10.1101/2022.05.23.493001

**Authors:** Bryce K. Allen, Meghana M. Kulkarni, Brian Chamberlain, Timothy Dwight, Cheryl Koh, Ramya Samant, Finith Jernigan, Jamie Rice, Dazhi Tan, Stella Li, Kristen Marino, Huang Huang, Evan Chiswick, Bethany Tesar, Sam Sparks, Zhixiong Lin, T. Dwight McGee, István Kolossváry, Charles Lin, Sharon Shechter, Holly Soutter, Cecilia Bastos, Mohammed Taimi, Sujen Lai, Alicia Petrin, Tracy Kane, Steven Swann, Humphrey Gardner, Christopher Winter, Woody Sherman

**Affiliations:** Silicon Therapeutics, 451 D Street, Boston, MA, USA, 02210

**Keywords:** STING, quantum mechanics, molecular dynamics, relative binding free energy simulations, artificial intelligence, innate immunity

## Abstract

The protein STING (stimulator of interferon genes) is a central regulator of the innate immune system and plays an important role in antitumor immunity by inducing the production of cytokines such as type I interferon (IFN). Activation of STING stems from the selective recognition of endogenous cyclic dinucleotides (CDNs) by the large, polar, and flexible binding site, thus posing challenges to the design of small molecule agonists with drug-like physicochemical properties. In this work we present the design of SNX281, a small molecule STING agonist that functions through a unique self-dimerizing mechanism in the STING binding site, where the ligand dimer approximates the size and shape of a cyclic dinucleotide while maintaining drug-like small molecule properties. SNX281 exhibits systemic exposure, STING-mediated cytokine release, strong induction of type I IFN, potent *in vivo* antitumor activity, durable immune memory, and single-dose tumor elimination in mouse models via a C*_max_*-driven pharmacologic response. Bespoke computational methods – a combination of quantum mechanics, molecular dynamics, binding free energy simulations, and artificial intelligence – were developed during the course of the project to design SNX281 by explicitly accounting for the unique self-dimerization mechanism and the large-scale conformational change of the STING protein upon activation. Over the course of the project, we explored millions of virtual molecules while synthesizing and testing only 208 molecules in the lab. This work highlights the value of a multifaceted computationally-driven approach anchored by methods tailored to address target-specific problems encountered along the project progression from initial hit to the clinic.

## 1 Introduction

Activation of the innate immune system through nucleic acid-sensing is a key mechanism for host defense against viruses and bacteria. Recent discoveries in the cyclic guanosine monophosphate-adenosine monophosphate synthase-stimulator of interferon genes (cGAS-STING) pathway have captured the attention of the pharmaceutical industry due to its role in innate immune signaling and the potential for pharmacologic modulation to treat autoimmune diseases Ahlers and Goodman (2016); Rui et al (2021), cancer Fuertes et al (2013); Wu et al (2020); Miao et al (2020), and viral infections Ahlers and Goodman (2016); Rodríguez-García et al (2018); Eaglesham et al (2019). STING is a mitochondria-associated membrane protein that is expressed in both immune and nonimmune cell types. When STING is activated through the binding of cyclic dinucleotides, including 2’,3’-cGAMP, which is produced by cGAS in response to cytosolic DNA Sun et al (2013), it undergoes a conformational change followed by oligomerization that induces type I interferon (IFN) and proinflammatory cytokines in a TANK-binding kinase (TBK1)–interferon regulatory factor 3 (IRF3)-dependent manner. The STING pathway is critically linked to antitumor immunity, including T cell activation in response to tumors, which requires STING expression in dendritic cells (DCs) and IFN-*β* expression via the STING pathway Fuertes (2011); Woo (2014).

X-ray crystallographic and cryogenic electron microscopy (Cryo-EM) studies have revealed that cGAMP binding to STING induces a pronounced conformational change from the ‘open’ inactive form of the ligand-free structure (Figure 1A) to a ‘closed’ active form that completely sequesters the bound ligand from solution in the buried binding site (Figure 1B) Shang et al (2012); Zhang et al (2019); Zhao et al (2019). Comparison of the STING apoprotein structure with the cGAMP-bound structure highlights a conformational transition that includes inward motion of the *α*1 helices, flap stabilization covering the cGAMP binding site, *β* sheet shift outward, and *α*4 modification (Figure 1B). The open (inactive) conformation adjusts to a closed (active) conformation when an agonist binds, where the central *α*2 helix from each monomer (residues 171-185) tilts by approximately 15*°*, and residues 226-241 in each monomer adopt a four-stranded *β*-sheet (not resolved by electron density in the open conformation). The distance between the ends of the central *α*2 helix is reduced by between *∼*18-20Å between the open and closed conformations upon activation. The thermodynamic equilibrium of these distinct conformational states suggests that stabilization of the open conformation could lead to inactivation of the cGAS-STING pathway, whereas stabilization of the closed conformation might lead to pathway activation.

**Fig. 1.**
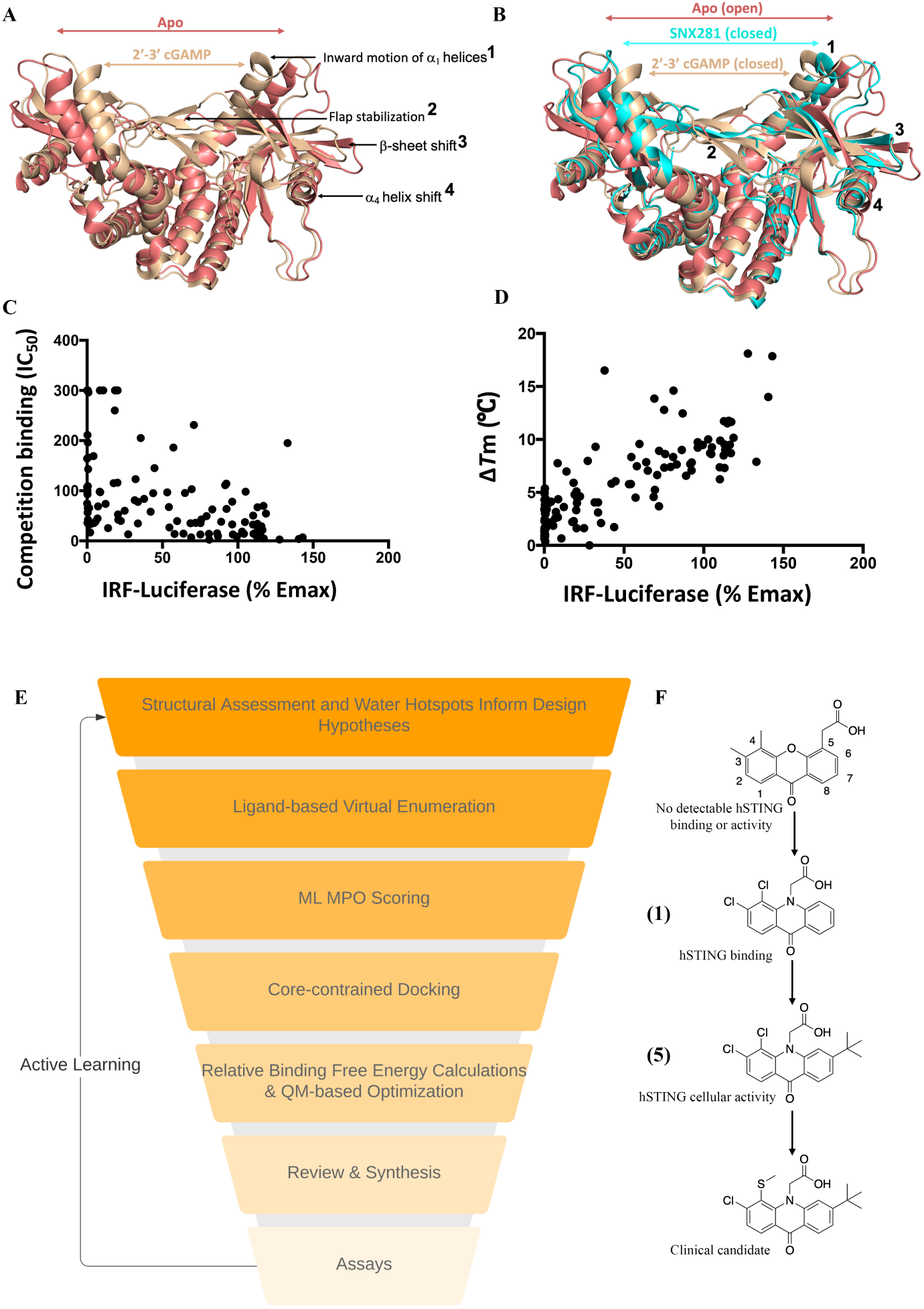
A) Overlay of apo (pink; PDB ID 4EMU) and 2’-3’-cGAMP bound (brown; PDB ID 4LOH) STING structures in ribbon representation, highlighting structural changes induced as STING changes from the open (inactive) to closed (active) conformation. Key structural features are labeled. B) Structure with SNX281 bound is additionally overlaid (cyan). C) Data demonstrating the disconnect between binding (IC_50_) and STING activation *in vitro*. D) Correlation is observed between STING activation and thermostabilization, as measured by differential scanning fluorimetry (DSF). E) Illustration of the virtual design workflow used in the optimization of SNX281. F) Summary of the progression from an initial molecule (DMXAA) with no human STING activity to the first screening hit possessing an acridone core **(1)**, intermediate molecule after 6-position optimization of an oxoacridinyl acetic acid hit **(5)**, and optimized clinical candidate SNX281.

cGAMP-triggered STING activation initiates downstream signaling by recruiting and activating TBK1, inducing the nuclear translocation of transcription factors IRF3 and NF-k*β*, culminating in the induction of type I IFN and proinflammatory cytokines such as tumor necrosis factor-alpha (TNF-*α*) Zhang et al (2019); Zhao et al (2019); Prabakaran et al (2018). Type I interferons are essential for robust anti-tumor immunity owing to their ability to stimulate T cell cross-priming, therefore improving the therapeutic potential of checkpoint blockade Fuertes (2011); Gkirtzimanaki et al (2018). Previous studies have demonstrated the therapeutic potential of STING agonism utilizing syngeneic mouse tumor models in which a cyclic dinucleotide (CDN) STING agonist exhibited marked anti-tumor activity when administered intratumorally, either alone or in combination with an inhibitor of programmed cell death protein (PD)-1 or PD-L1 Martin et al (2019). However, the poor pharmacokinetic properties of CDN-based STING agonists limited administration to direct intratumoral injection Wang et al (2020); Novotná et al (2019).

The feasibility of treating tumors with a systemically dosed STING agonist was first demonstrated in mouse models two decades ago with the xanthone derivative DMXAA Baguley and Ching (2002); Gao et al (2013b); Weiss et al (2017). However, due to structural differences between the human and murine variants of the STING protein Hwang et al (2019), those results did not translate to human efficacy, owing to the lack of sufficient binding and conformational stabilization of the human STING protein (hSTING). Subsequent efforts to utilize DMXAA as a starting point to design a hSTING agonist have been unsuccessful Baguley and Ching (2002); Gao et al (2013b); Hwang et al (2019). More recent studies have presented small-molecule STING agonists suitable for systemic administration that efficiently bind to hSTING, which could be applicable to a broad range of malignancies Hong et al (2021); Sintim et al (2019), with some entering clinical development. The efforts outlined above highlight the unique challenges associated with the discovery of a clinically-relevant systemic STING agonist, which include: (1) design of a molecule with desirable drug properties in the context of a large polar binding site suitable for endogenous CDNs, (2) engineering a requisite level of binding with a precise conformational modulation that translates across the relevant STING isoforms, and (3) a mechanism of action that activates STING both rapidly but briefly, in order to minimize potential on-target toxicity that may result from sustained NFk*β* activation. Given the above factors, our aim was to create a systemically available C*_max_*-driven agonist that would stabilize the appropriate conformation of the hSTING dimer and lead to downstream activation (as opposed to maximizing binding affinity). In this work, we describe the use of physics-based simulations and artificial intelligence to rationally engineer SNX281, a systemically available small molecule STING agonist that activates hSTING through a unique dimerization mechanism and is currently in Phase I clinical trials, both as a monotherapy and in combination with a PD-1 inhibitor (Figure 1E,F).

## 2 Results

The unique nature of the STING protein – a homodimer that undergoes CDN-induced activation via a large-scale conformational change – necessitated a novel design approach and the development of bespoke computational methods to predict binding affinity, conformational stabilization, and drug properties (Figure 1E). Inspired by the understanding that two molecules of DMXAA bind to a single mouse STING (mSTING) homodimer Gao et al (2013b), we sought to apply a computational strategy in order to engineer chemical matter capable of binding and activating human STING (hSTING) to produce IFN-*β* that could also bind and activate in the relevant efficacy (mouse) and toxicology (rat and monkey) STING species to maximize our chances of successfully translating preclinical animal data to the clinic. We hypothesized that the dimeric binding of DMXAA in mSTING is possible in part due to the large binding site with two-fold rotational (C2) symmetry, which enables two copies of DMXAA (*∼*300 Dalton molecular weight) to fill the CDN binding pocket, approximately matching the size of a CDN (*∼*700 Da; see Figure 2A,B).

**Fig. 2.**
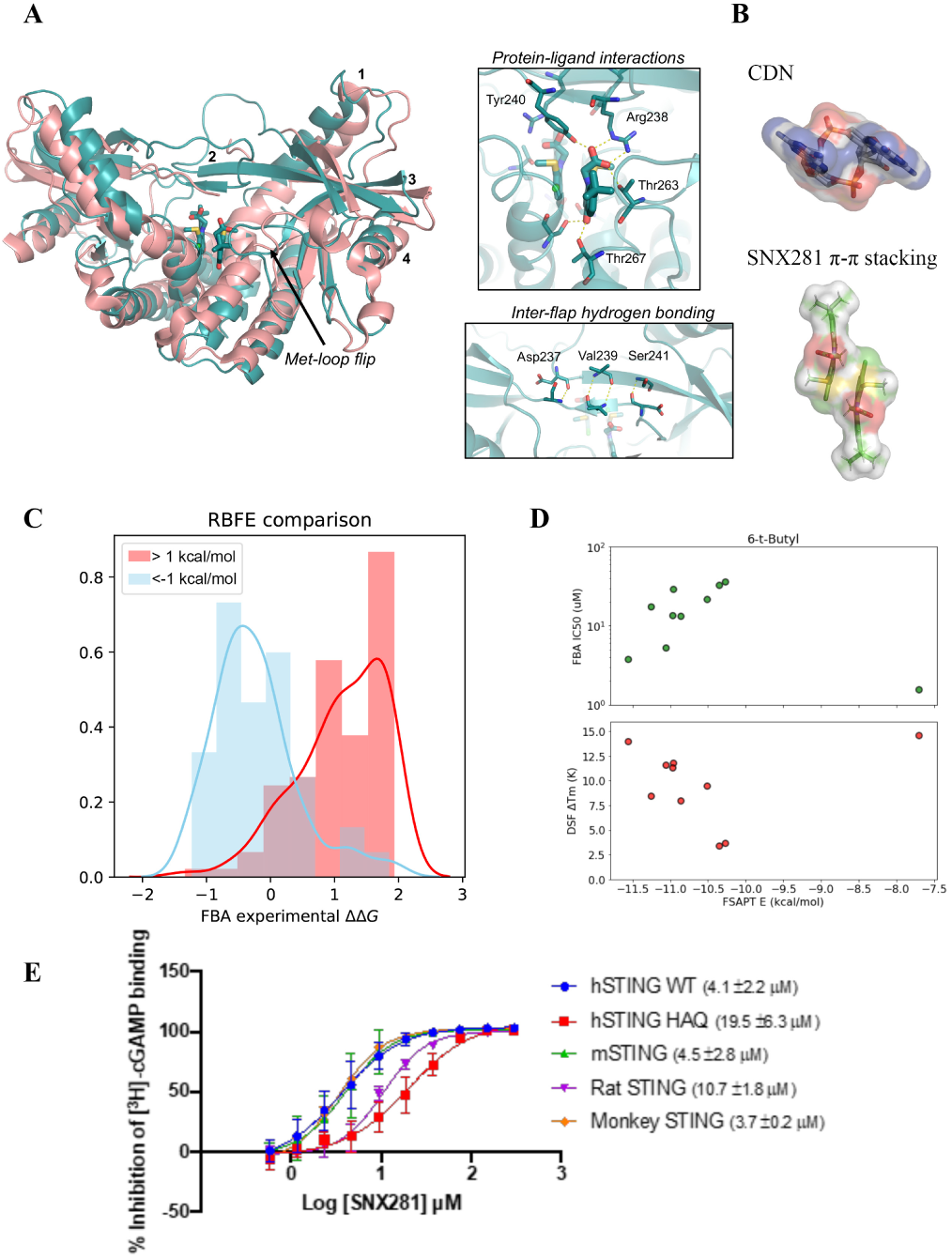
A) Key interactions induced by SNX281 binding to hSTING including met-loop flipping, protein-ligand interactions, and inter-flap hydrogen bonding. B) Shape comparison of endogenous CDNs and SNX281, which interact with each other in the binding site through *π−π* stacking. Surface color scheme shows (N, O, S, and C) as blue, red, yellow, and green, respectively. C) Predictive power of the bespoke relative binding free energy (RBFE) workflow developed in this work, which assesses multiple thermodynamic pathways for the dimeric nature of binding. D) Correlation between the *π*-*π* interaction energy from F-SAPT with the experimental IC_50_ from a fluorescent binding assay (FBA; top) and differential scanning fluorimetry (DSF; bottom) for a subset of compounds containing a t-butyl moiety at the 6-position (6-t-butyl). E) SNX281 displaces [3H]-cGAMP from hSTING in a homogeneous filtration-based competition assay. Plot shows % inhibition of [3H] - cGAMP binding to human wild type (hSTING WT) and HAQ variant (hSTING HAQ), mouse (mSTING), rat and monkey STING by SNX281. Error bars indicate SD. IC_50_ (*µ*M) of inhibition is shown in parentheses.

### 2.1 SNX281 Design Strategy

We developed a multi-pronged computational strategy to design small molecules that optimized binding and conformational stabilization of hSTING while maintaining desirable drug-like properties. Iterative cycles of molecular design, predictive simulations, chemical synthesis, biological assays, and data analysis (design, predict, make, test, analyze; DPMTA) were conducted until we arrived at a molecule with the desired properties (SNX281). Methods for computational predictions were developed throughout the project based on the nature of the challenges at hand. Specifically, we: (1) identified binding hotspots through water thermodynamic calculations, which led to initial binding (Supplementary Figure 1), (2) stabilized face-to-face *π − π* stacking within the ligand dimer using quantum mechanics, which led to initial activation, and (3) leveraged molecular dynamics (MD) simulations to design molecules that enhanced activation by stabilizing a specific conformation of hSTING. Throughout the design process, ideas were generated through traditional medicinal chemistry approaches and augmented with virtual molecules enumerated using an active learning framework. Design ideas were evaluated by multi-parameter optimization (MPO) using computationally predicted properties that were used to progress the chemical series toward the target product profile (TPP), ultimately resulting in SNX281.

The computational tools developed and implemented in this project are briefly described below:

#### Identification of binding hotspots from water thermodynamics

MD simulations of the hSTING protein surrounded by thousands of water molecules were analyzed using the Grid Inhomogeneous Solvation Theory (GIST) methodology Ramsey et al (2016), where water entropy is computed based on spatial and orientational correlation functions using frames from the MD trajectory and enthalpy is computed from pairwise interaction energies. The resulting water thermodynamics were used to identify regions of unfavorable water molecules (hotspots) that revealed regions of the hSTING binding pocket where significant boosts in binding affinity could be gained upon displacement by an appropriate ligand functional group Breiten et al (2013); Beuming et al (2012).

#### Virtual enumeration of design ideas

Molecules were enumerated based on synthetic accessibility rules derived from publicly available template sets Hartenfeller et al (2011). Common reactions included alkylations, condensations (amides and sulfonamides), palladium-catalyzed couplings, and protecting group manipulations. Between 10,000-100,000 virtual molecules were generated in each iteration of the DPMTA cycle.

#### Multi-parameter optimization (MPO)

Based on the desired target product profile (TPP), as described above, we defined an MPO objective function that removed molecules with undesirable predicted properties and enriched the dataset in molecules that could achieve the desired profile. In each iteration, over 90% of the virtual molecules were eliminated. The remaining virtual molecules were subjected to more computationally expensive methods to prioritize for chemical synthesis and testing (see below).

#### Lead prioritization

Molecules that passed the MPO filter described above were prioritized for synthesis and experimental evaluation using a suite of computational tools that we developed throughout the course of the project, based on insights gained from molecular simulations, x-ray crystal structures, and iterative DPMTA feedback:

1. **QM-based dimer interaction energy** - Quantum mechanics (QM)-based functional group symmetry-adapted perturbation theory (F-SAPT) was used to optimize face-to-face *π−π* stacking within the dimeric molecules Parrish et al (2018). We chose to implement a QM-based approach for the dimerization interactions because traditional molecular mechanics force fields do not accurately account for interactions where polarization plays a significant role, such as the *π − π* interaction between the two molecules in the hSTING binding site. F-SAPT precisely predicts the quantum mechanical interaction energy (including polarization) between user-defined functional groups (substitutions on the acridone core in this case).
2. **Alchemical binding affinity prediction** - Alchemical relative binding free energy (RBFE) calculations leverage a closed thermodynamic cycle to gain computational efficiency by ”perturbing” one molecule into another. Such calculations are more computationally efficient than directly simulating a binding event for each molecule. RBFE is most appropriate when molecules of interest have significant parts in common, as in our hit-to-lead and lead optimization efforts of the early oxoacridinyl acetic acid hits Cournia et al (2017). The unique dimerization mechanism of SNX281 and preceding acridinyl molecules in the project necessitated the development of a novel dual-perturbation workflow that properly considered the two molecules in the binding site interacting with each other and the protein (Dual-RBFE) Song et al (2019); Lee et al (2020a); Cournia et al (2017). In short, we perturbed candidate molecules along two pathways within the thermodynamic cycle (Figure 2), combining the results to determine the binding affinity and confidence estimates.
3. **Active Learning** - In each design iteration, candidate molecules were enumerated, evaluated, and progressed toward the desired TPP while leveraging the uncertainty of experimental data and *in silico* predictions, including calculations of solubility, permeability, clearance, cytochrome P450 (CYP) inhibition, hERG inhibition, binding affinity, and cellular activity.

### 2.2 Structure-Activity Relationship (SAR) for SNX281 Chemical Series

Computational enumeration and *in silico* assessment of small functional group libraries at each chemistry handle on the acridone core identified only a few promising positions where activity could be improved (positions 3, 4, and 6 of the acridone core; see Figure 1F for numbering). However, early structure-activity relationships (SAR) for modifications at these positions highlighted the challenge of designing molecules with improved affinity, conformational activation, and drug-like properties. A conformational change in the ‘met-loop’ (residues 262-272), as determined by x-ray crystallography, proved to be a distinguishing feature for 6-position modifications that improved binding (Figure 2A). Insights regarding the the nature of 6-position modifications came from the identification of a high-energy water patch that could only be accessed from the 6-position (Supplementary Figure 1). Table 2 lists several examples of SAR (*in silico* and *in vitro*) that illustrate the early challenges to improve potency. Although Dual-RBFE calculations supported the replacement of methyl with various substitutions, these modifications neither induced ‘met-loop’ preferential flipping nor led to improvements in activity, with the exception of the t-butyl derivative **5** (Table 2).

**Table 1:**
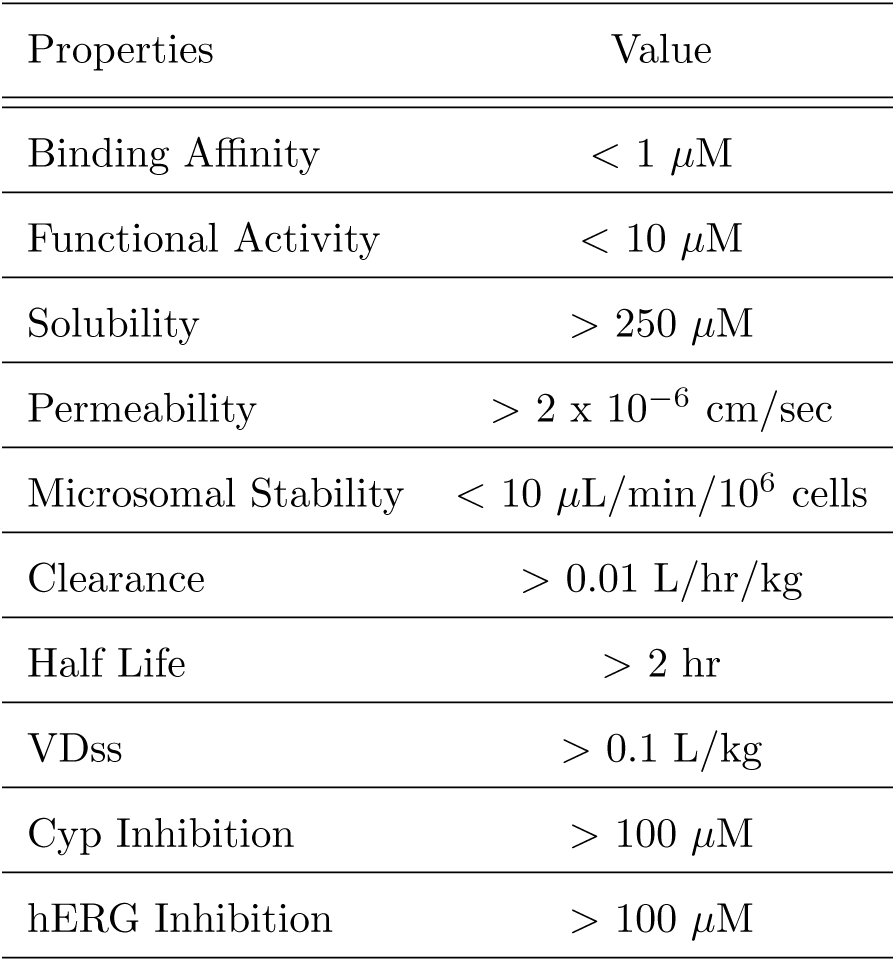
Multiparameter optimization profile considered during the design of SNX281.

**Table 2:** SAR for exemplary 6-position (R_1_) analogs of the acridone core. FBA = fluorescence-based assay; RBFE = relative binding free energy, as computed by thermodynamic integration; GP = Gaussian process, a supervised learning approach used to solve regression and probabilistic classification problems.

| Compound | $R_1$ | $\Delta pK_d$ | $\Delta pK_d$ | $\Delta pK_d$ |
| --- | --- | --- | --- | --- |
|  |  | FBA | RBFE | GP |
| **1** → **2** | | $0.54 \pm 0.23$ | $0.75 \pm 0.16$ | $-0.41 \pm 0.51$ |
| **1** → **3** | | $-0.39 \pm 0.12$ | $-0.87 \pm 0.18$ | $0.57 \pm 0.45$ |
| **1** → **4** | | $1.15 \pm 0.29$ | $1.58 \pm 0.26$ | $-0.26 \pm 0.34$ |
| **1** → **5** | | $0.96 \pm 0.22$ | $1.21 \pm 0.22$ | $0.32 \pm 0.32$ |

Explorations at the 4-position showed minimal predictive capabilities from Dual-RBFE alone, while F-SAPT augmentation of the Dual-RBFE calculations resulted in better ranking of the designed molecules (Table 3). Consistent with previous observations regarding substituent effects on face-to-face aromatic interactions Meyer et al (2003), dimer decomposed association energies correlated well with the electron richness of the 4-position substituents as well as steric interaction/repulsion across the dimer interface (Table 3). For example, the electron-rich ethoxy (**12**) has a more favorable predicted dimer association energy while maintaining predicted binding energy via RBFE, explaining its improved overall binding activity. At the same time, naphthyl (**13**) and cyclopropyl (**14**), being more electron-poor, exhibited less self-interaction energy, as predicted by F-SAPT, which may contribute to their weaker STING binding and activation. Encouraged by these calculations, additional designs were evaluated to simultaneously explore subtle variations in the 4- and 6-positions to enhance ligand-ligand and protein-ligand interactions.

**Table 3:** SAR for selected analogues exploring 4 position.

| Compound | $R_2$ | $\Delta pK_d$ | $\Delta pK_d$ | $\Delta pK_d$ |
| --- | --- | --- | --- | --- |
|  |  | FBA | RBFE | GP |
| **5** → **6** | | $0.09 \pm 0.15$ | $0.05 \pm 0.18$ | $0.92 \pm 0.43$ |
| **5** → **7** | | $-0.64 \pm 0.12$ | $-0.24 \pm 0.28$ | $-0.63 \pm 0.61$ |
| **5** → **8** | | $1.3 \pm 0.07$ | $1.18 \pm 0.18$ | $0.32 \pm 0.47$ |
| **5** → **9** | | $1.07 \pm 0.17$ | $1.54 \pm 0.34$ | $-0.42 \pm 0.52$ |

In total across design cycles, 2,039,481 molecules were explored using reaction-based molecular enumeration complemented with ideas from medicinal chemists and molecular modelers. This pool of molecules, which were filtered with MPO ADMET predictions as described above, reduced the number to 54,829 molecules that were computationally explored with Dual-RBFE and a subset (1,293) with F-SAPT. A total of 208 molecules were synthesized for *in vitro* characterization across the duration of the project, culminating in our clinical candidate (SNX281). Experimental results were consistent with our prospective predictions (see Figure 2C,D) and supported the hypothesis that hSTING activation by our chemical series depends on a delicate balance of ligand-ligand and ligand-protein interactions, coupled with the stabilization of a specific conformation of hSTING leading to activation (Figure 2C).

### 2.3 *In Vitro* Characterization of SNX281

On-target binding activity of SNX281 was evaluated in homogeneous filtration-based competition and protein thermal shift assays using recombinant human, mouse, rat and monkey STING proteins. SNX281 competes with and inhibits the binding of radiolabeled cyclic dinucleotide (CDN) 2’ 3’-cGAMP (3H-cGAMP), to the C-terminal domain of human STING with an IC_50_ of 4.1 *±* 2.2 *∼ µ*M. Importantly, binding activity of SNX281 is maintained across all species tested (mouse, rat and monkey) as well the clinically relevant HAQ haplotype of human STING (Figure 2E). Structural analyses of the STING protein show that the symmetric dimer undergoes a conformational transition from an inactive ‘open’ state to an active ‘closed’ conformation upon interaction with agonists Kranzusch (2015); Gao et al (2013b); Zhang et al (2013); **?**.

Binding of the bacterial cyclic di-GMP and the endogenous 2’3’-cGAMP do not equally induce this conformational change and result in varying degrees of stabilization of the active closed state. Analysis of the cellular activity of STING bound to different cyclic dinucleotides and the small molecule ligand DMXAA suggest that ligands that better stabilize the ”active” state result in more robust activation of downstream signaling and IFN-*β* induction. Consistent with these observations, thermal stabilization of the symmetric STING dimer by ligand binding was a better predictor of cellular activity than binding affinity alone. SNX281 stabilizes the thermal unfolding of STING with a ΔTm of 12.2*^◦^*C (Table 4). While the binding affinity of SNX281 for STING is less than that of its endogenous CDN agonist, thermal stabilization of the STING protein, a hallmark of *bona fide* ligand binding, by SNX281 is superior to that by the CDN (ΔTm of 12.2 versus 10.9*^◦^*C respectively) (Table 4).

**Table 4:** *ΔTm(◦C)* is the difference in melting temperature between apo and ligand bound STING and measures the thermal stabilization of recombinant human, mouse, rat and monkey STING proteins upon SNX281 binding.

|  | Human |  | Mouse | Rat | Monkey |
| --- | --- | --- | --- | --- | --- |
| Ligand | WT | HAQ | WT | WT | WT |
| SNX281 | 12.2 ± 1.4 | 13.9 ± 1.4 | 27.1 ± 4.2 | 12.3 | 11.6 |
| cGAMP | 10.9 ± 0.5 | 22.2 ± 0.7 | 25.2 ± 0.2 | 11.3 | 15.4 |

Direct binding of SNX281 to STING was confirmed by X-ray crystallography. The 2.3*∼*^°^A cocrystal structure of SNX281 bound to hSTING (residues 151 to 343) is shown in Figure 2A. Crystallographic studies of early hits from the SNX281 chemical series confirmed the dimeric binding, where the acridone core makes a face-to-face *π − π*-stacking interaction with its partner (Figure 2A). The two molecules of SNX281 recapitulate the key interactions made by the CDNs, such as the salt-bridge with Arg238 to induce the ‘closed’ conformation of the flap (Figure 2B). SNX281 binding induces conformational transitions that are known to be required for downstream signaling including the inward motion of the *α*1 helices, flap stabilization, *β*-sheet shift and *α*4 helix shift (Figure 1A,B). SNX281 makes predominantly hydrophobic interactions with human STING, punctuated by several anchoring polar contacts. The carboxylate of each molecule of SNX281 forms a salt bridge with Arg238 and a hydrogen bond with Thr263 of each chain of the STING monomers. In addition, the carbonyl on each acridone core forms a hydrogen bond with Thr267 of each chain (Figure 2A). The 3 and 4-position substituents on the acridone core increase thermodynamic stabilization through the *π − π* stacking, as predicted by F-SAPT, further driving the conformational equilibrium toward the ‘closed’ state (Figure 2D). The two ligand monomers form a substantial dimer interface within the active site, burying approximately 230*∼*^°^A^2^ of solvent-accessible surface area (SASA; Fig S2), emphasizing the importance of ligand-ligand interactions in the observed binding mode.

SNX281 is highly selective, exhibiting no significant binding against a panel of 44 receptor, transporter, ion channel, and enzyme assays when tested at 10 *µ*M (Table S2, CEREP results). Consistent with its small size and drug-like properties, SNX281 also exhibited higher permeability than CDNs in an *in vitro* passive permeability assay across Caco-2 cell monolayers (apparent permeability = 6.6 10*^−^*^6^cm/sec*^−^*^1^ for SNX281 versus undetected for CDNs).

### 2.4 SNX281 is a STING agonist in cells

Following activation by ligand binding, STING translocates from the ER to the ER-Golgi intermediate complex (ERGIC) where it associates with TBK1, which in turn mediates the phosphorylation and activation of the transcription factors IRF3 and NFk*β*. Once activated IRF3 and NFk*β* translocate into the nucleus to drive the expression of type I IFNs and the down-stream cytokine/chemokine cascade Chen et al (2016). To determine the functional consequence of binding to STING, HEK293T expressing STING (HAQ variant) cells were treated with SNX281 and analyzed for induction of phosphorylation and changes in subcellular localization of STING, TBK1 and IRF3 proteins as well as downstream induction of cytokines. Treatment of HEK293T cells expressing STING with SNX281 induced STING translocation from the ER to peri-nuclear foci, TBK1 phosphorylation and co-localization with STING, and IRF3 cytoplasm-to-nuclear translocation in a dose dependent manner (Figure 3A). In THP-1 cells, treatment with SNX281 resulted in rapid (*<*30 minutes) phosphorylation of STING (S366), TBK1 (S172) and IRF3 (S396) proteins (Figure 3B) and caused dose dependent induction of IFN-*β* with an EC50 of 6.6*µ*M (Figure 3C) consistent with the observed binding potency. Relative to DMSO vehicle control, SNX281 induced higher levels of phosphorylation than either the same concentration of ADU-S100 (Adu CDN) or 10 times the concentration of 2’3’-cGAMP (Figure 3B). In addition to IFN-*β* and consistent with the activation of the NFk*β* pathway downstream of STING, SNX281 also induced the release of the proinflammatory cytokines Interleukin 6 (IL-6) and tumor necrosis factor alpha (TNF-*α*) (Figure 3C) in treated cells.

**Fig. 3.**
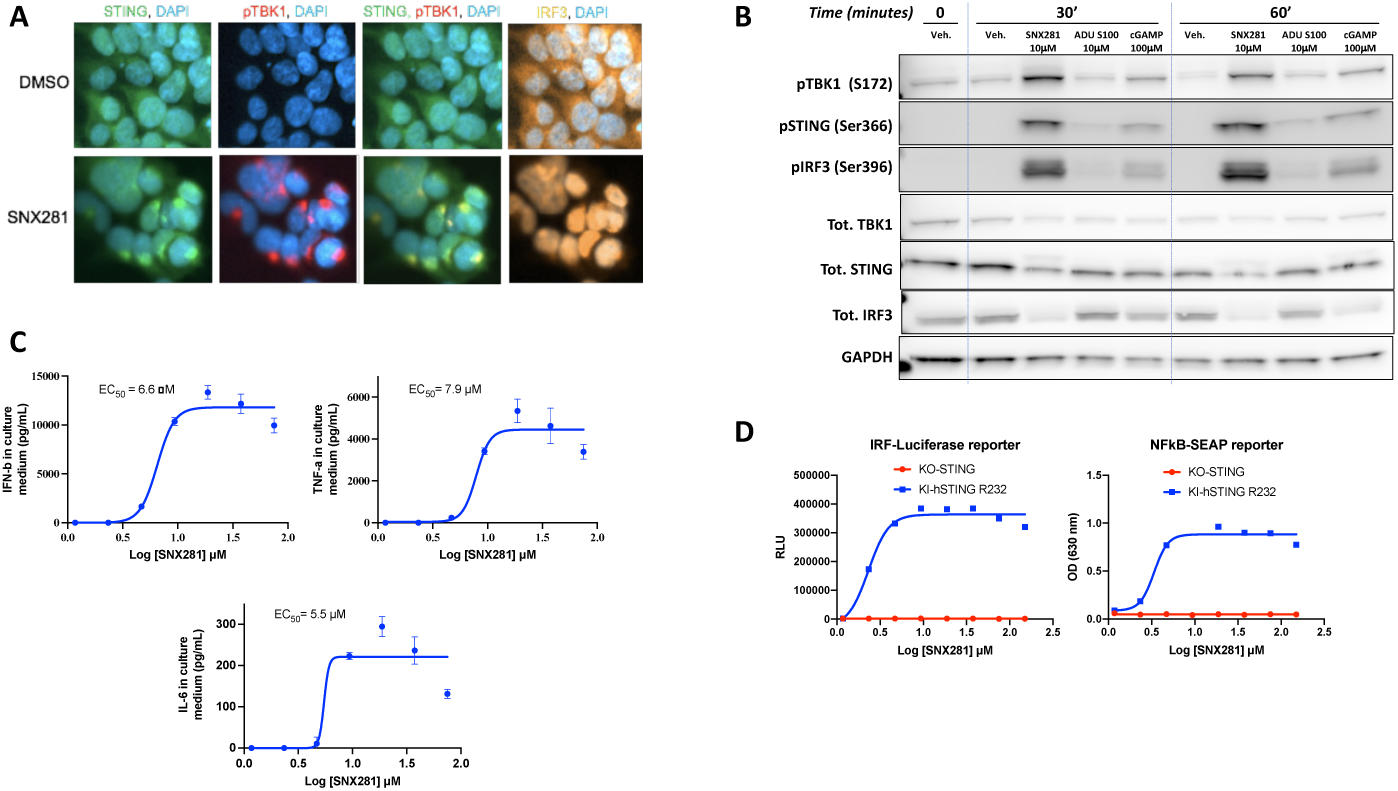
Confirmation of SNX281 as a STING agonist. A) HEK293T cells stably expressing HAQ human STING protein were treated with 25 *µ*M SNX281 or DMSO for 1 hour. Cells were fixed, stained with antibodies for STING, phospho-TBK1(Ser172) and IRF3 proteins prior to imaging. SNX281 induces STING translocation from ER to peri-nuclear foci, TBK1 phosphorylation and co-localization with STING, and IRF3 cytoplasm-to-nuclear translocation. B) Lysates from THP-1 cells treated with 10*µ*M SNX281, 10*µ*M ADU-S100, 100*µ*M 2’-3’ cGAMP or DMSO (Vehicle control) for 30 minutes were analyzed by western blotting for the expression of STING, phospho-STING (Ser366), phospho-IRF3 (Ser396), and phospho-TBK1(Ser172). Lysates from THP-1 cells treated with varying concentrations of SNX281 were collected after 0.5, 1, 1.5 and 2-hour exposure and analyzed by western blotting with antibodies specific to STING and phospho-STING (Ser366). To quantify the level of STING phosphorylation induced by SNX281 the peak areas of the chemiluminescent signals corresponding to phospho-STING (Ser366) and total STING protein were determined using the Compass by SW software. Data points correspond to peak area (High Dynamic Range 4.0) corresponding to phospho-STING (Ser366) expressed as a percentage of the peak area (High Dynamic Range 4.0) corresponding to total STING for treated samples relative to DMSO controls. C) THP-1 cells were stimulated with SNX281 for 6 hours, and the levels of secreted IFN-*β*, TNF-*α* and IL-6 in the cell culture supernatant were measured by ELISA. SNX281 induced the secretion of IFN-*β*, TNF-*α*, and IL-6 in a dose-dependent manner with EC_50_ for induction of each cytokine shown above. D) The effect of SNX281 on the IRF and NFk*β* induction in THP-1 cells was evaluated using luciferase and SEAP reporters respectively.

Treatment of THP1-Dual reporter cell lines expressing either wild-type hSTING or the common HAQ allele with SNX281, resulted in the induction of IRF3-luciferase and to a lesser extent, the NFkB-SEAP reporter gene (Table S1). The cellular EC50 for IRF3-reporter induction by SNX281 is 1.1 *±*0.63*µ*M (WT) and 1.7*±*0.53*µ*M (HAQ) respectively and is *>*10-fold more potent than the endogenous CDN agonist 2’ 3’-cGAMP (Figure 3D), reflecting poor membrane permeability of the CDN despite its high binding affinity. SNX281 treatment of STING-deficient (KO) cells does not induce reporter gene expression, demonstrating that SNX281, like 2’ 3’-cGAMP, acts specifically via STING (Figure 3D).

### 2.5 Human allele specificity and cross species activity of SNX281

Single nucleotide polymorphisms (SNPs) in the human STING gene affect its ability to be activated by canonical CDN agonists Gao et al (2013a,c); Diner et al (2013). Five polymorphic variants of human STING (WT, REF, HAQ, AQ and Q) that vary at amino acid positions 71, 230, 232 and 293 have been identified Jin et al (2011); Yi et al (2013). To test the ability of SNX281 to activate these common human STING variants each of the full-length proteins was stably expressed in HEK293T cells carrying ISRE-SEAP as a measure of IRF3 activity, and IFN-*β*-luciferase reporter genes Diner et al (2013). HEK293T cells are known to lack endogenous STING expression Burdette and Vance (2013) and do not respond to CDNs in the absence of ectopic expression of the STING protein. SNX281 potently activated all common variants of human STING expressed in HEK293T cells, resulting in rapid phosphorylation of STING (Supplementary Figure S2) and in the induction of reporter genes several fold higher than that induced by 2’3’-cGAMP CDN (Figure 4A). Stimulation of peripheral blood mononuclear cells (PBMCs) from a panel of human donors carrying different STING alleles (hSTING-WT/WT, hSTING-WT/REF, hSTING-WT/HAQ, hSTING-REF/REF, and hSTING-HAQ/REF) with SNX281 *in vitro* induced IFN-*β* expression. The EC_50_ of IFN-*β* induction across these donors ranged from 3.5-9.9*µ*M (Figure 4B and Supplemental Table 1). A summary of cytokine induction from SNX281 stimulated PBMCs from a panel of 11 human donors is noted in Supplemental Table 1.

**Fig. 4.**
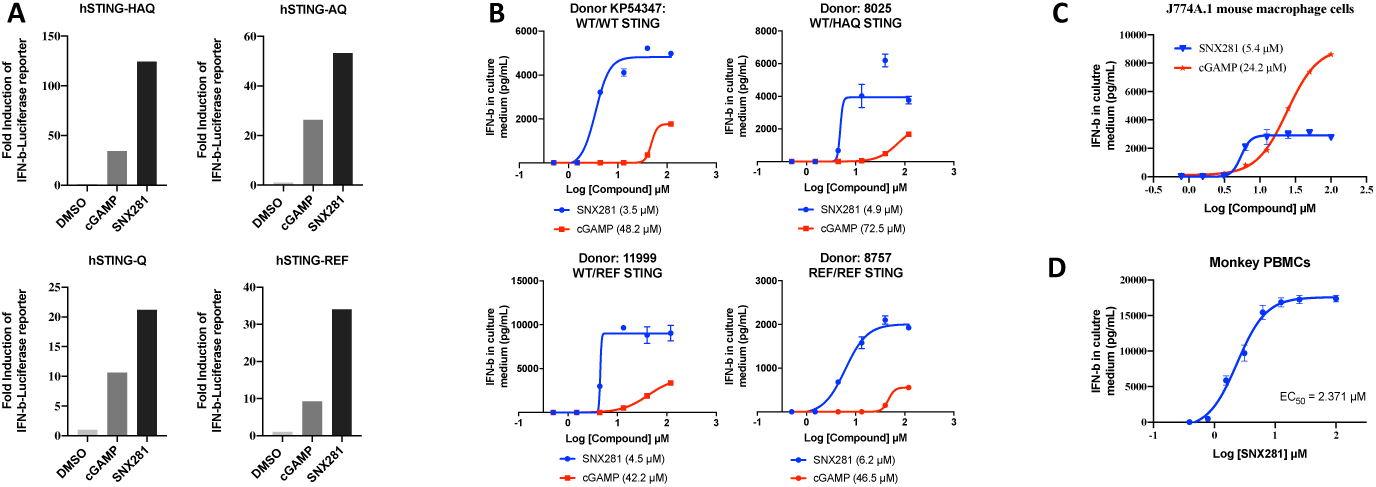
Human allele specificity and cross species activity of SNX281. (A) HEK293T cells stably expressing each of the full-length human STING variants and carrying ISRE-SEAP and IFN-*β*-luciferase reporter genes were treated with DMSO, CDN (2’3’-cGAMP; 25*µ*M), or SNX281 (20788; 25*µ*M). Fold induction of IFN-*β*-luciferase reporter relative to control (DMSO) treated samples is plotted. (B) IFN-*β* secretion from the PBMCs of 4 donors upon stimulation over a concentration range of SNX281 or cGAMP. STING variant is indicated the graph for each donor with corresponding EC50 values of SNX281 and cGAMP indicated below each graph in brackets; similarly the IFN-*β* concentration-response to SNX281 and cGAMP of (C) J774A.1 murine macrophage cells (EC_50_s in brakets) and in (D) IFN-*β* secretion from cynomolgus monkey PBMCs in response to SNX281 (EC_50_=2.37*µ*M).

Administration of SNX281 to cells leads to dose-dependent induction of IRF3- and NFk*β*-driven reporters (Figure S3) as well as IFN-*β* (Figure 4C) secretion in mouse macrophage-like cell line (J774A.1) with an EC_50_ of 5.4*µ*M compared to 24.2*µ*M for cGAMP. To determine activity of SNX281 to rat and cynomolgus monkey STING protein each full-length protein was stably expressed in HEK293T cells carrying the ISRE-SEAP (measure of IRF3 activity) and IFN-*β*-luciferase reporter genes Diner et al (2013). Treatment of cells with SNX281 resulted in the induction of reporter genes several fold higher than that induced by 2’3’-cGAMP (Figure S4). Stimulation of PBMCs from cynomolgus monkey with SNX281 *in vitro* resulted in a dose dependent increase in the secretion of IFN-*β* detected by ELISA with an EC_50_ of 2.4*µ*M (Figure 4D).

### 2.6 Pharmacokinetic profile and pharmacodynamic activity of systemic SNX281: A *hit and run* mechanism

As prolonged induction of inflammatory cytokine signaling may be harmful to the host, STING activation and downstream signaling is tightly regulated to ensure robust and timely response against infection while minimizing risk associated with prolonged immune response. Endogenous STING protein is rapidly degraded after activation as it translocates from the ER to vesicles via the endolysosomal pathway Gonugunta et al (2017). We hypothesized that therapeutically efficacious and safe systemic STING stimulation could be achieved by an agonist with rapid distribution, strong activation, and a short half-life, thereby mimicking physiological STING activity dynamics.

To better understand the mechanistic pharmacology of SNX281 activation we evaluated the induction of the signature cytokine IFN-*β* in mouse and cynomolgus monkey over a 24-hour period after SNX281 administration, with a focus on characterizing the concentration-response relationship and time dependence of induction. C57BL/6 mice (n=4) were treated with a single dose of vehicle (control group) or 10, 25, 35, and 45 mg/kg SNX281 administered intravenously (i.v. bolus). Pharmacokinetic analysis indicated SNX281 has a short half-life of 2.33 hours (Figure 5A). IFN-*β*, the pharmacodynamic marker of STING activation, was induced in a dose-dependent manner at 3 hours post-dose (Figure 5B) in mice. Blood samples were collected at 3- and 6-hours post-dose for determination of plasma IFN-*β* levels using a qualified ELISA. IFN-*β* was undetectable at all time points in the plasma of vehicle treated control animals. As shown in Figure 5B at 3-hours, the plasma level of IFN-*β* in mice and increased in a dose-dependent manner in response to SNX281. In SNX281 treated mice, at all doses the levels of IFN-*β* were the highest at the 3-hour time point, relatively reduced at 6-hours (Supplementary Figure S5) and were undetectable at 24-hours (Supplementary Figure S5). These data indicate that SNX281 rapidly but transiently induces a potent type I IFN response *in vivo*. To translate more closely to humans, we characterized the pharmacological activity in cynomolgus monkeys. This indicated that the level of IFNb induction was correlated with the C*_max_* concentration of SNX281 (Figure 5C).

**Fig. 5.**
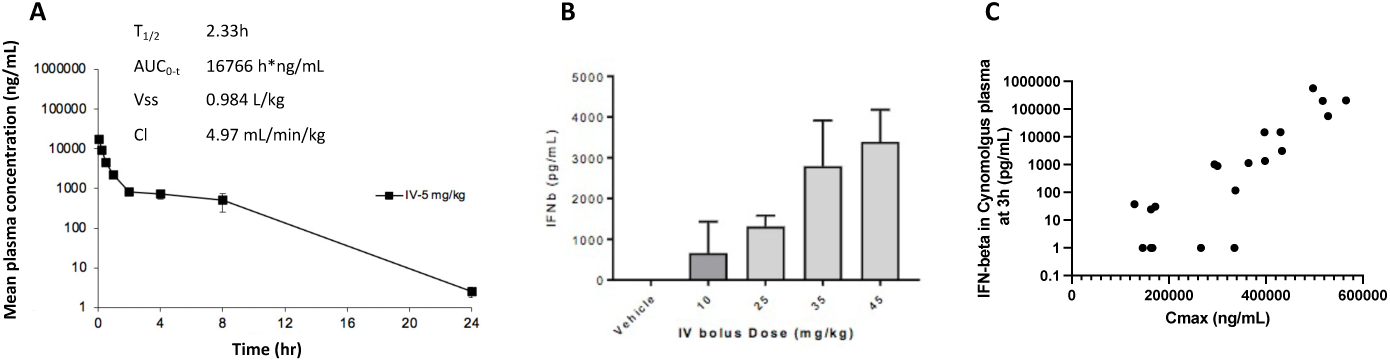
Pharmacokinetic profile and pharmacodynamic activity of systemic administration of SNX281 in mouse and cynomolgus monkey. (A) Plot of the mean plasma concentration vs. time after a single i.v. dose of 5 mg/kg. Estimated PK parameters are shown. (B) Plasma IFN-*β* concentrations induced with increasing dose levels of SNX281.(C) Relationship between plasma IFN-*β* levels and C*_max_* in cynomologus monkeys after treatment with SNX281.

### 2.7 SNX281 monotherapy demonstrates robust dose-dependant anti-tumor activity

The anti-tumor activity of SNX281 was tested in syngeneic tumors in mice with established CT26 (colorectal) tumors; n=8 per group) received a single i.v. bolus dose of vehicle (Group 1), 35 mg/kg (Group 2) or 45 mg/kg (Group 3) of SNX281. In addition to a single i.v. administration at the high dose levels, mice were also treated with multiple i.v. injections at lower dose levels of SNX281 as: 12.5 mg/kg (Group 4) on D1/4/7, 25 mg/kg (Group 5) on D1/4/7, and 25 mg/kg (Group 6) b.i.d (2 hours apart) on D1. All mice in Groups 2 (35 mg/kg) and 3 (45 mg/kg) were completely cured of tumors after receiving a single i.v. dose of SNX281. Dose-dependent anti-tumor response was observed with 12.5 (D1/4/7), 25 (D1/4/7), 25 b.i.d (2 hours apart on D1) and 35 (D1) mg/kg resulting in 0/8, 1/8, 3/8 and 8/8 complete responses (CR) respectively (Figure 6A).

**Fig. 6.**
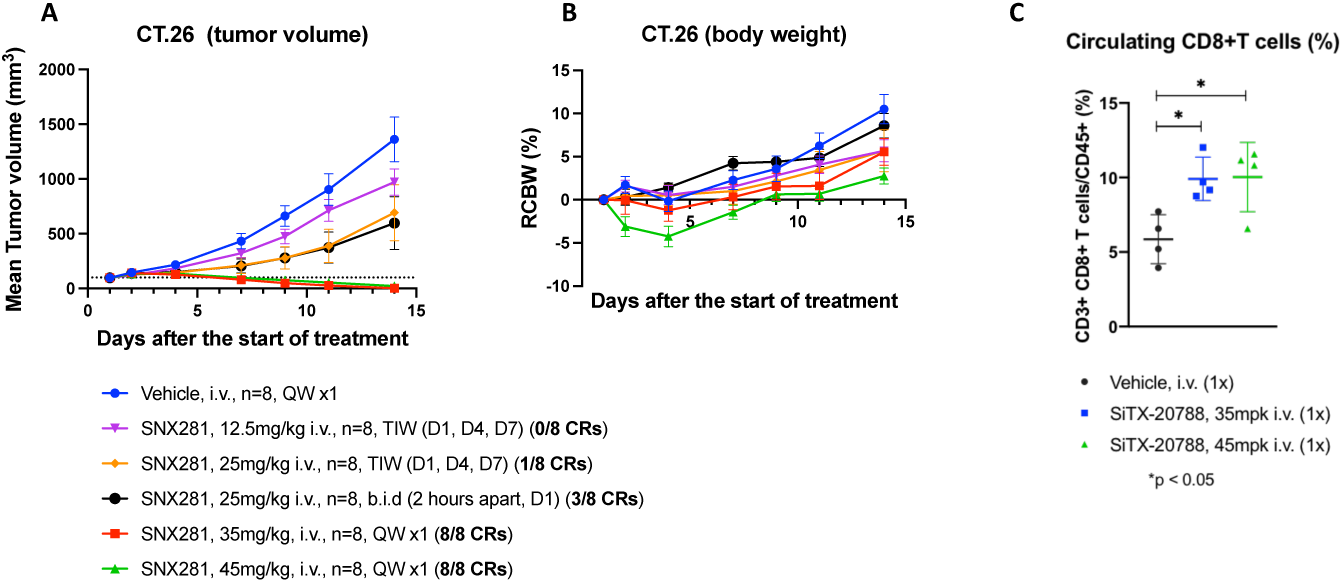
SNX281 monotherapy demonstrates robust dose-dependent, long term anti-tumor activity. A) and B) BALB/c mice were inoculated subcutaneously on the right flank with 3 x 105 CT26.WT cells (n = 8/group). When tumor volumes were 98 mm^3^, mice received Vehicle (i.v., QWx1); 35 or 45 mg/kg SNX281 (i.v., QWx1); 12.5 mg/kg SNX281 (i.v., TIW D1, D4, D7); 25 mg/kg SNX281 (i.v., TIW D1, D4, D7 or 25 mg/kg SNX281 (i.v., b.i.d on D1, 2 hours apart). A) Tumor measurements were taken every two days. Data points represent group mean and error bars represent standard error of the mean (SEM). B) Body weights were measured every two days. Shown is percent Relative Change of Body Weight (%RCBW) for each mouse calculated according to the following formula 1. BW*_TreatmentDayN_* is the average body weight of a treatment group on a given day, BW*_TreatmentDay_*_0_ is the average body weight of the treatment group on the first day of treatment. Data points represent percent group mean change in body weight. Error bars represent standard error of the mean (SEM).

The observation that tumor regression was induced in the CT26 model by a single i.v. dose of 45 mg/kg, but not by two doses of 25 mg/kg separated by two hours provides evidence for the requirement for a C*_max_* over a threshold level for optimal dimerization and STING activation (Figure 6A). There was no significant loss of body weight observed in mice across all treatment groups (Figure 6B) demonstrating that systemic STING activation by SNX281 was well tolerated. On D17, whole blood was collected from mice in Groups 1/2/3 for flow cytometric analysis using a lymphocyte cell panel comprised of CD45, CD3, CD4, CD8, CD25, Ki67 and H-2L*^d^*-AH1 surface markers. A *∼*2-fold increase (*P <* 0.05) in the number of circulating CD8+ T cells (Figure 6C) and a *∼*3-fold increase in tumor antigen-specific, AH1+ CD8+ T cells was observed in SNX281 treated mice compared to the vehicle treated group (Supplementary Figure S6).

### 2.8 SNX281 demonstrates long-term anti-tumor activity

To investigate whether SNX281 could elicit durable tumor-specific T-cell responses, mice that were cured of their primary tumor were re-challenged with either the same or different tumor cells. Mice cured of primary tumors (Groups 2 and 3) as well as 10 näıve controls were housed under similar conditions for 3 months and enrolled into a tumor re-challenge study on Day 85 (D85) post the initial treatment with SNX281. Four out of eight cured mice in each of Groups 2 and 3 were subcutaneously (s.c.) inoculated on the left flank with 3x10^5^ CT26.WT cells while the remaining (4/8) cured mice were inoculated with unrelated A20 lymphoma cells (5x10^6^ cells/mouse). 5 näıve mice were also enrolled as control group for each cell line. Both cell lines (CT26.WT and A20) developed tumors normally in näıve mice. In contrast, mice that were previously cured of primary tumors by SNX281 (35 and 45 mg/kg) were significantly protected from re-challenge with CT26.WT cells and 4/4 mice remained tumor free for over a month (Figure 6D). Additionally, one of four cured mice from each dose group was also protected from re-challenge with the unrelated A20 lymphoma cells. Thus, a single i.v. administration of the efficacious doses of SNX281 imparts protection from tumor re-challenge demonstrating the induction of durable immunity.

### 2.9 Anti-tumor activity of SNX281 requires the host immune system

In order to determine the contribution of STING activation in host immune cells versus tumor cells, we tested the anti-tumor activity of SNX281 in CT26.WT tumor-bearing, immune compromised NOG mice. The NOG (NOD/Shi-*scid* /IL-2R*δ^null^*) mouse is a severely immunodeficient mouse, developed by Central Institute for Experimental Animals (CIEA) in 2000 (https://www.ciea.or.jp/en/laboratory_animal/nog.html). NOG (NOD/Shi-*scid* /IL-2R*δ^null^*) mice have multiple immunodeficiencies including lack of functional T and B cells, lack of NK cells, dendritic cell, and macrophage dysfunction and defects in complement hemolytic activity. Treatment of CT26.WT tumor bearing NOG mice (average TV *∼*100 mm^3^) with 35 and 45 mg/kg of SNX281 (n=8 mice/group) did not result in any significant anti-tumor activity as had been previously observed in immunocompetent wild-type mice (Figure 7A and B). This demonstrated that an intact host immune system is required to mediate the anti-tumor activity of SNX281.

**Fig. 7.**
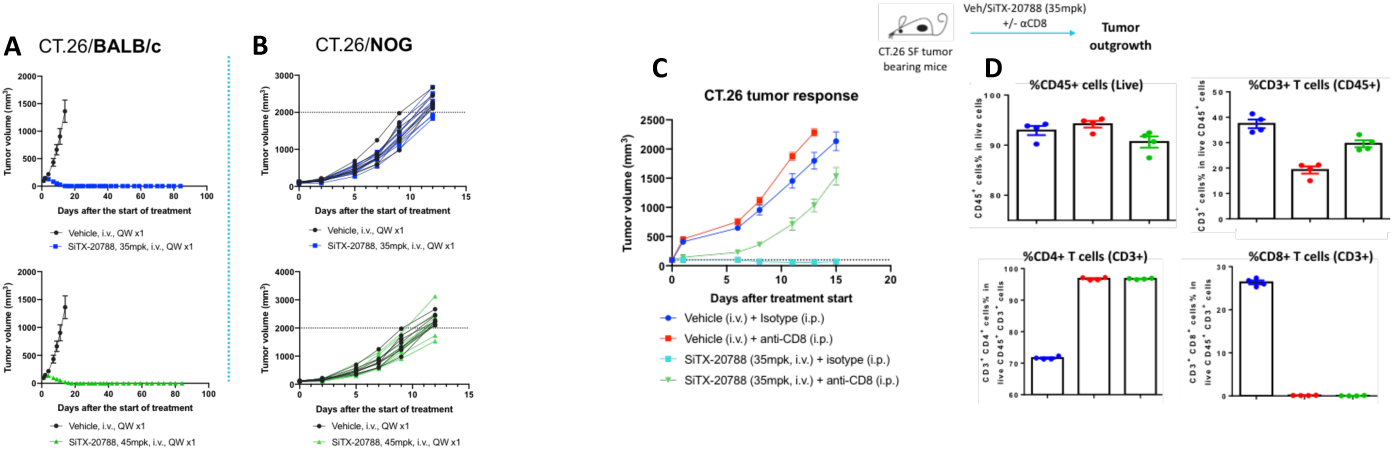
Anti-tumor activity of SNX281 requires the host immune system. A) BALB/c mice were inoculated subcutaneously on the right flank with 3 x 10^5^ CT26.WT cells (n = 8/group). When tumor volumes were 98 mm^3^, mice received Vehicle (i.v., QWx1); 35 (top) or 45 (bottom) mg/kg SNX281 (i.v., QWx1). All mice were completely cured of tumors after receiving a single i.v. dose of 35 or 35 mg/kg SNX281. B) Severely immunodeficient mouse NOG (NOD/Shi-*scid* /IL-2R*δ_null_*) mice were inoculated subcutaneously with 3x105 CT26.WT cells on the right flank for tumor development. When tumor volumes were *∼*107 mm^3^, mice received Vehicle (i.v., QWx1); 35 (top) or 45 (bottom) mg/kg SNX281 (i.v., QWx1). The tumor sizes were measured three times per week till the end of study on Day 12. C) BALB/c mice bearing single right flank CT26.WT tumors were i.p. administered 25 mg/kg isotype control or anti-mouse CD8*α* antibody on days -2, -1, 1, 5, 10 and 15; with one i.v. injection of vehicle or 35 mg/kg SNX281 on Day 0. Tumor volumes were measured three times per week and shown is mean tumor growth over time (n=8 mice/group). Error bars represent standard error of the mean (SEM). D, Flow cytometric analysis from peripheral whole blood on Day 4 confirmed *>*95% depletion of CD8 + T cell population in mice treated with anti-CD8*α* antibody

To determine the role of SNX281-induced CD8+ T cells in tumor control, mice bearing single (right) flank CT26.WT tumors were depleted of CD8+ T cells before and after i.v. bolus dose of SNX281. Specifically, tumor bearing mice received a single i.v. dose of vehicle or 35 or 45 mg/kg SNX281 when average tumor volumes were *∼*97 mm^3^ (defined as D0). Additionally, mice in vehicle- and SNX281-treated groups were intraperitoneal (i.p.) administered either 25 mg/kg of Isotype IgG control or anti-mouse CD8a antibody on Day -2, -1, 1, 5, 10 and 15 (n=8 mice/group). The depletion of CD8+ T cells accelerated (31.3% increase in mean tumor volume; *P <* 0.0001) the growth of tumors in the vehicle control group. CD8+ T cell depletion also prevented the control of tumors treated with 35 mg/kg of SNX281 (48.3% difference in mean tumor reduction in isotype versus anti-CD8a -treated mice, *P <* 0.0001) (Figure 7C). Flow cytometry data from peripheral whole blood on Day 4 confirmed the profound depletion of CD8+ T cells in mice treated with anti-mouse CD8a antibody compared with IgG isotype control treated mice (Figure 7D). Thus, maximal tumor control depended on host CD8+ T cells, confirming the involvement of an adaptive immune component in SNX281 mediated anti-tumor activity, although some tumor control was observed even in the absence of T cells.

### 2.10 SNX281 Enhances Anti-tumor Activity in Combination with Checkpoint Inhibitors

The ability of SNX281 to enhance T-cell–mediated antitumor immune responses in combination with checkpoint inhibitors was tested in multiple tumor models that are resistant to anti-PD-1 alone. BALB/c mice bearing single (right) flank CT26.WT tumors (average TV *∼*100^3^) were treated with vehicle or SNX281 (25 mg/kg) via i.v. bolus administration once a week for three weeks (qwk x3). The vehicle and SNX281 treated mice were also adminis-tered via i.p injections either rat IgG2a isotype control antibody or anti-mouse PD-1 (RMP1-14) BIW for three weeks (n=8 mice/group). As has been previously observed for CT26 tumor model, no significant effect on tumor growth was observed with anti-PD-1 alone. Significant TGI (66% TGI; *P <* 0.001) was observed in mice treated with 25 mg/kg SNX281 which was further enhanced by combining with anti-PD-1 (88% TGI; *P <* 0.0001) (Figure 8A). In addition to enhanced anti-tumor activity the combination of SNX281 and anti-PD-1 had improved survival benefit (Figure 8B). Parallel studies were conducted to characterize changes in T cell responses caused by SNX281 (25 mg/kg), anti-PD-1 or combination treatment. On Day 7 post treatment initiation, mice treated with either SNX281 alone or the combination of SNX281 and anti-PD-1 antibody showed an increase in circulating tumor-antigen specific AH1+ CD8+ T cells (Figure 8C). IFN-g ELISpot analysis of peripheral PBMCs on day 14 post treatment initiation showed a significant (*P <* 0.001) enhancement in the functionality of circulating T cells in mice treated with SNX281 alone or in combination with anti-PD-1 antibody (Figure 8D).

**Fig. 8.**
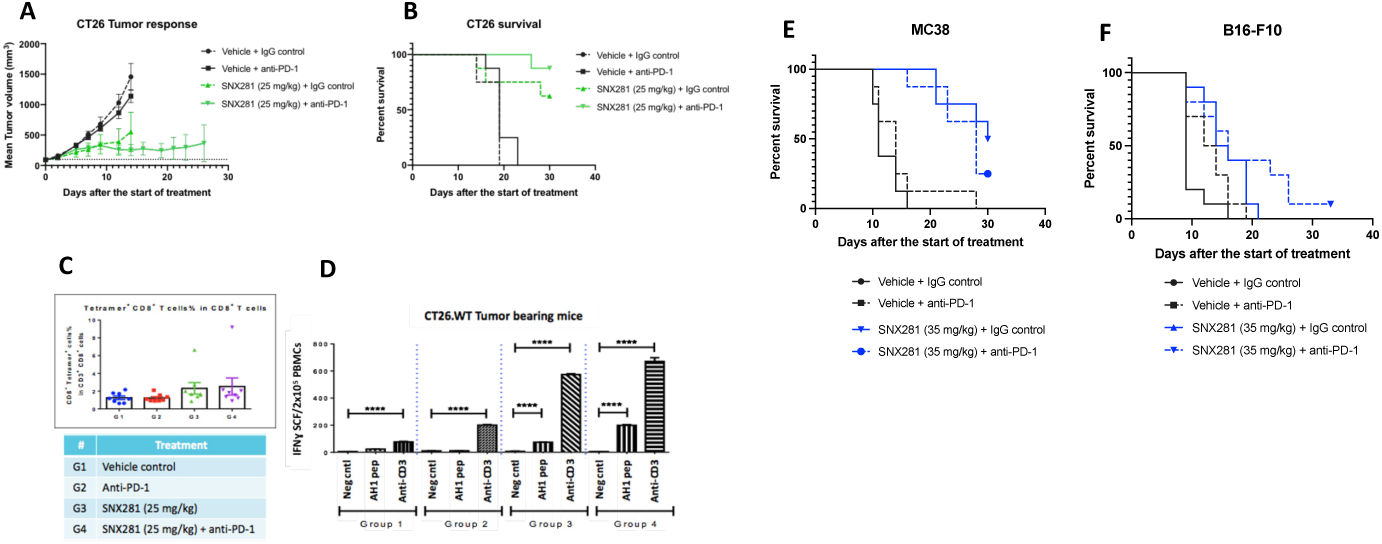
SNX281 enhances anti-tumor activity in combination with checkpoint inhibitors. A) BALB/c mice bearing single right flank CT26.WT tumors (*∼*95mm^3^) were injected via the intraperitoneal (i.p.) route with isotype control or anti-mouse PD-1 antibody 5 mg/kg on days 0, 3 and 10 mg/kg on days 7, 10, 14 and 17; with i.v. injection of vehicle or 25 mg/kg SNX281 on days 0, 7 and 14. Tumor volumes were measured three times per week and shown is mean tumor growth over time (n=8 mice/group). Error bars represent standard error of the mean (SEM). B) For each animal, survival time was defined as the time from treatment initiation to termination. Percent survival curve was plotted using the Kaplan-Meier method. C) and D) BALB/c mice (n=8 mice/group) bearing single right flank CT26.WT tumors (*∼*95mm^3^) were injected via the i.p. route with isotype control or anti-mouse PD-1 antibody 5 mg/kg on days 0, 3 and 10 mg/kg on days 7, 10, 14 and 17; with i.v. injection of vehicle or 25 mg/kg SNX281 on days 0, 7 and 14. C, PBMCs from mice in all four groups was collected 7 days post SNX281 dose initiation and assessed for the frequency of CD8+ H-2Ld-AH1 Tetramer+ (AH1+) cells by Flow cytometry. D) 14 days post SNX281 dose initiation, peripheral blood mononuclear cells (PBMCs) from mice in all four groups was collected and stimulated with media (Neg Ctrl), AH1 peptide or anti-CD3 antibody (positive control) and assessed by IFN-*γ* enzyme-linked immunospot (ELISpot).

C57BL/6 mice bearing single (right) flank B16-F10 tumors (average TV *∼*60 mm^3^) were treated with vehicle or SNX281 (35, 45 mg/kg) via i.v. bolus administration once a week for two weeks (qwk x2). The vehicle and SNX281 treated mice were also administered either 10 mg/kg rat IgG2a isotype control antibody or anti-mouse PD-1 (RMP1-14) i.p. BIW for two weeks (n=8 mice/group). In this poorly immunogenic melanoma tumor model, treatment with SNX281 (35 or 45 mg/kg) alone significantly inhibited the growth of tumors relative to vehicle treated control group (53% and 61% TGI respectively on D7; *P <* 0.0001) (Figure 8E). Survival gains were demonstrated with either SNX281 (*P <* 0.001) or checkpoint inhibitor alone compared to vehicle + isotype, while the combination of 45 mg/kg SNX281 and anti-PD-1 demonstrated significantly improved survival benefit over anti-PD-1 monotherapy alone (P=0.001) (Figure 8F). 3/10 mice treated with 45 mg/kg SNX281 alone were found dead on Day 9 (two days after the second dose), as were 3/10 treated with 45 mg/kg SNX281 and anti-PD-1; thus the combination with anti-PD-1 did not result in additional toxicity.

Thus, combining STING-dependent T cell priming induced by SNX281 with anti-PD-1 resulted in robust anti-tumor activity and significant survival benefit in multiple tumor models. Importantly, the anti-tumor effect of SNX281 in tumors with distinct histology as well as in different host genetic backgrounds indicates the broad therapeutic potential of SNX281.

### 2.11 C*_max_*-Driven Pharmacology of SNX281

The assembly of activated STING dimers into oligomers and subsequent translocation and signaling is a complex process. While activation of the TBK1/IRF3 axis and activation of NFk*β* both occur downstream of STING activation, the results of the SNX281 exposure with post exposure wash in THP1 cells suggest that a brief period of SNX281 exposure induced activation of the IRF pathway without activating NFk*β* (Figure 9). This pharmacological effect may result from the kinetics of concentration driven SNX281 dimerization in STING, and is therapeutically advantageous as it may favor induction of IFN-*β* release over inflammatory cytokine release when dosed systemically by a brief IV infusion or bolus. The *in vivo* observation that tumor regression was induced in the CT26 model by a single IV dose of 45 mg/kg, but not by two doses of 25 mg/kg separated by two hours provides evidence for the requirement for a C*_max_* over a threshold level for optimal dimerization and STING induction (Figure 9).

**Fig. 9.**
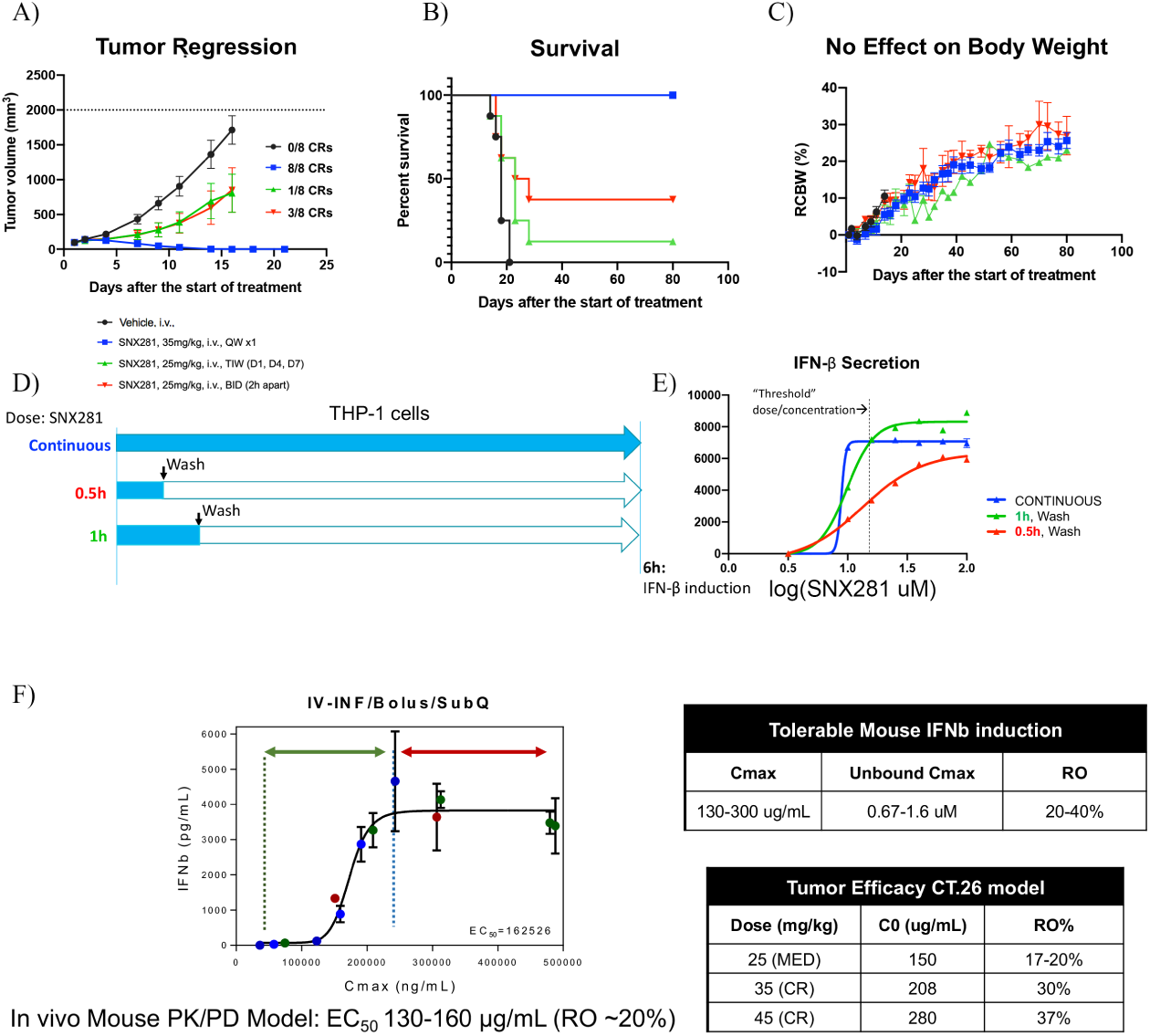
Efficacy is driven by a short exposure over an optimal threshold. A-C) Tumor regression, survival, and effect on body weight is shown for SNX281. D,E) One hour exposure over the threshold concentration is sufficient to drive durable IFN-b response in cells. F) Exposure-response analysis in mouse model reveals concentration-dependent for efficacious type I IFN induction.

## 3 Discussion

We report the design of SNX281, a systemically available, non-nucleotide–based STING agonist with demonstrated *in vivo* anti-tumor activity. The unique challenges presented by the STING protein necessitated the development of bespoke computational tools to account for the non-covalent self-dimerization of compounds in the binding site of STING and requisite conformational stabilization of the activated state of STING. Using a suite of predictive computational tools developed specifically to solve the challenges presented by STING, we designed a systemically available small molecule STING agonist with drug-like properties that could fill the large, polar active site of STING and stabilize the active conformation through a unique self-dimerizing binding mode. When used as a single agent in mice, systemically administered SNX281 induces tumor regression with durable anti-tumor immunity and is well tolerated. In mouse tumor models that are poorly responsive to PD-1 blockade, combinations of SNX281 with an anti–PD-1 immune checkpoint inhibitor are superior to monotherapy in inhibiting tumor growth and prolonging survival. These observations support the concept that induction of host adaptive anti-tumor immunity via pharmacological activation of STING can overcome tumor resistance to checkpoint inhibitors.

First-generation CDN-based STING agonists that entered clinical trials were dosed by intratumor injection. A recent study by Sivick et al. highlighted the challenges associated with attaining optimum drug levels by direct intratumoral dosing, a critical requirement to balance the immunogenic and cell-ablative effect of STING activation Sivick et al (2018). In contrast, IV administration of SNX281 and the corresponding pharmacokinetic profile allow for a broader range of tumors and fine-tuning of STING agonism in various tissues *in vivo*. SNX281 is systemically available and is expected exhibit higher cellular potency in an acidified tumor microenvironment (versus normal tissue) because it is a weak acid. Previous work on a different chemical series of weak acids suggested that preferential activation of STING in tumors substantially contributes to the observed favorable *in vivo* anti-tumor activity and tolerability profilePan et al (2020).

## 4 Methods

### 4.1 Biochemical and biophysical methods

Compound binding affinity to STING was measured using a competition assay with tritiated 2’,3’-cGAMP (3H-cGAMP), for with wild type (WT) or the HAQ variant using a homogeneous, in-solution filter-binding assay (FBA) (Li et al. 2014). FBA assay conditions were 250nM 6xHis-SUMO STING (aa140-379) protein, 25nM 3H-cGAMP, PBS buffer (ph7.4), 1 hour. Inhibition 3H-cGAMP was measured with co-incubation of SNX281 over a concentration range from 1mM to 0.5*µ*M using a Microbeta radiation counter for detection. Differential Scanning fluorimetry (DSF) was used to monitor the thermal denaturation profile STING proteins in the absence or presence of a concentration range of compound (Niesen et al. 2007). DSF assay conditions were 5*µ*M purified 6xHis-SUMO STING (aa140-379), 1x SYPRO Orange Dye, SNX281 over a concentration range (30nM-100*µ*M) in buffer (30*µ*M Hepes, ph7.5 with 100*µ*M NaCL). Tm was measured using a ramp from 25*°*C to 95C on ViiA7 qPCR machine (ABI) over 30min.

Protein Expression Purification Plasmid of 6His-SUMO-hSTING construct was transformed into E.coli BL21(DE3) cells. 50 mL small culture was grown in LB medium at 37*°*C for overnight. Larger scale bacterial cultures were induced with IPTG at OD600 reached 0.6-0.8 and then was attained in incubator at 20*°*C for another 16-20 hrs. Cell pellet were resuspended with lysis buffer (30 mM Tris pH 8.0, 150mM NaCl, 10% glycerol, 0.05% tween-20, 0.5 mM TCEP) + SUMO Express Protease, lysed with sonication, and then centrifuged at 30,000 g at 4*°*C for 30 mins. The supernatant was loaded onto a Ni-NTA affinity column, and protein was eluted with buffer: 30 mM Tris pH 8.0, 150 mM NaCl, 100 mM Imidazole, 0.5 mM TCEP. The eluted protein was loaded onto HiTrap Desalting column and was exchanged into buffer desalt (30 mM Tris pH 8.0, 150 mM NaCl, 1 mM TCEP). The N-terminal 6His-SUMO tag was then cleaved by incubating the eluted protein with TEV protease at 4*°*C overnight. After TEV cleavage, the hSTING protein was further purified by SEC (size-exclusion chromatography) using HiPrep 26/60 Sephacryl S-200 HR column equilibrated with buffer containing (30 mM Tris pH 8.0, 150 mM NaCl, 3% glycerol, 0.5 mM TCEP).

Crystallization, data collection, and structural determination Initial crystallization screening was done at 20*°*C using sitting-drop vapor-diffusion method. The protein concentration for hSTING crystallization was 8.5 mg/ml. Optimization of the hits were carried out in 3*µ*L hanging-drop vapor diffusion method. The crystals of compound + hSTING were grown in 0.1M Magnesium format dihydrate, 11% PEG 3,350, 5% Glycerol. Cryo-solution for all protein crystals was 20% glycerol plus reservoir solution. HKL2000 was used to process the X-ray diffraction data. Molecular replacement using APO structure of hSTING as searching template was performed by using PHENIX software. All the three-dimensional structural images of protein and ligand were made in PyMOL.

### 4.2 Cellular Assays

High content imaging of STING agonism: HEK293T stably expressing multiple isoforms of human STING protein were treated with 25*∼µ*M SNX281 or DMSO for 1 hour. Cells were fixed, stained with antibodies for STING, phospho-TBK1(Ser172) and IRF3 proteins. Agonist-induced translocation from ER-peri-nuclear foci was imaged and quantified using EnVision Reader (PerkinElmer, Waltham, MA). Cells were fixed after exposure to compound for 1 hour and stained for with antibodies against STING, phos-TBK1 and IRF3. Lysates from THP-1 cells treated with 10*∼µ*M SNX281, 10*∼µ*M ADU-S100, 100*∼µ*M 2’-3’ cGAMP or DMSO (Vehicle control) for 30 minutes were analyzed by western blotting for the expression of STING, phospho-STING (Ser366), phospho-IRF3 (Ser396), and phospho-TBK1(Ser172). Lysates from THP-1 cells treated with varying concentrations of SNX281 were collected after 0.5, 1, 1.5 and 2-hour exposure and analyzed by western blotting with antibodies specific to STING and phospho-STING (Ser366).

Activation of human sting polymorphic variants in engineered HEK293T cell lines HEK293T-Dual-Null cells, lacking STING (Invivogen). 293 Dual-Null cells stably express an ISRE (Interferon stimulated response element)-inducible SEAP (secreted embryonic alkaline phosphatase) reporter construct upon IRF3 binding. These cells are also engineered to express Lucia luciferase under control of the IFN-*β* promoter. This reporter cell line was used to generate stable cell lines expressing four common polymorphic variants of human STING (HAQ, AQ, Q and REF) that vary at amino acid positions 71, 230, 232 and 293 from the WT allele. Additionally, 293-Dual cells stably expressing rat and monkey STING proteins were generated in order to assess the cross-species activity. All lines were confirmed for expression of full-length STING by Western blotting and the identity of various allelic variants by Sanger sequencing of cDNA. IFN-*β* induction was measured after 20 hours of compound incubation at concentrations listed in Figure 4. Luciferase activity was measured using Quanti-Luc reagent with cell supernatant and luminescence measured on a Spectramax i3 spectrophotometer.

IFN inflammatory cytokine secretion Cryopreserved human PBMCs from a cohort of healthy donors of varying STING genotypes, including the wild type, HAQ and REF alleles, were stimulated with SNX281 or cGAMP for 6 hours. The levels of secreted IFN-*β*, TNF-*α*, IL-6 and IL-1*β* in the cell culture supernatant were quantified by enzyme linked immunosorbent assay (ELISA). PBMCs were isolated from donor leukopaks using a standard procedure with Ficoll centrifugation. Donor PBMCs were genotyped for STING variants using Sanger sequencing. Cytokine secretion was measured from PBMC supernatants (48 well plates; 2 x 10^6^ cells/well) after treatment with compound for 6 hours. Human IFN-*β* TNF-*α*, IL-6 and IL-1*β* were quantified using the DuoSet ELISA Development System (RD Systems) performed as per manufacturer’s instructions. Activation of STING in murine cells was tested using the J774A.1 macrophage-like cell line. After a 6-hour incubation with compound the amount of secreted IFN-*β* was measured in the cell culture by ELISA according to manufacturers instructions (mouse IFN-*β* DuoSet ELISA, RD Systems). STING-mediated induction IFN-*β* in PBMCs from cynomolgus monkey (IQ Biosciences; donor 5032, Lot P19D2206) was tested after treatment for 6h using an ELISA (VeriKine Cynomolgus IFNb ELISA) with cell supernatants.

### 4.3 *In vivo* Methods

151 BALB/c mice were inoculated subcutaneously on the right flank with CT26.WT cells for tumor development. Seven days after tumor cell inoculation, 96 mice with tumor volumes ranging from 80-120*∼mm*^3^ (average tumor volume around 98*∼mm*^3^) were selected and assigned into 12 groups using stratified randomization with 8 mice per group based upon their tumor volumes. Compound treatment was started from the day of randomization (defined as D1). Group 1 was treated with 10% DMAC + 90% 50 mM NMG/water, i.v. on D1, Group 2 was treated with SNX281 35mg/kg i.v. on D1, Group 3 was treated with SNX281 45mg/kg i.v. on D1, Group 4 was treated with SNX281 12.5mg/kg i.v. on D1/4/7, Group 5 was treated with SNX281 25mg/kg i.v. on D1/4/7, Group 6 was treated with SNX281 25mg/kg i.v. BID on D1 (2 hours apart). Tumor sizes and body weights were measured three times per week and mice with tumors exceeding 2000*∼*mm^3^ were euthanized. On D17, 0.1*∼*mL whole blood was collected for flow cytometry (FCM) analysis from the remaining mice in Group1 and from 4 out of 8 mice in Groups 2 and 3.

Studies assessing the host immune contribution were performed in immunodeficient mice and mice depleted of CD8+ T-cells. For the former, 36 severely immunodeficient NOG mice (NOD/Shi-scid/IL-2Rnull)NOG mice were inoculated subcutaneously with CT26.WT cells on the right flank for tumor development. Six days after tumor cell inoculation, 24 mice with tumor volumes ranging from 79-132*∼*mm^3^ (average tumor volume 107 mm^3^) were selected and assigned into 3 groups using stratified randomization (the randomization start day was defined as D0) with 8 mice in each group based upon their tumor volumes. Group 1 was treated with Vehicle, i.v., QW x1; Group 2 was treated with SNX281, 35 mg/kg, i.v., QW x1 and Group 3 was treated with SNX281, 45 mg/kg, i.v., QW x1, respectively. The tumor sizes were measured three times per week till the end of study on D12. Survival was monitored and mice with tumor volumes exceeding 2000*∼mm*^3^ considered to be the humane endpoint, were euthanized according to protocol. To test the role of CD8+ cells, 84 BALB/c mice were inoculated subcutaneously with CT26.WT cells on the right flank for tumor development. Mice were divided into two clusters (40 mice in cluster 1 and 44 mice in cluster 2). Cluster 1 was pretreated with Isotype IgG control antibody (MAb rat IgG2*β*, BioXcell) and cluster 2 was pretreated with anti-mouse CD8*α* antibody (MAb anti-mouse CD8*α* antibody, CAT BE0061, BioXcell) by intraperitoneal (i.p.) administration on D-2, D-1 before randomization (defined as D0). Nine days after tumor inoculation, 24 mice from each pretreated cluster with tumor volumes ranging from 83-114*∼*mm^3^ (average tumor volume 97*∼*mm^3^) were selected and assigned into 3 groups using stratified randomization with 8 mice in each group based upon their tumor volumes. Groups 1/3/5 were selected from isotype pretreated mice (cluster 1) and Groups 2/4/6 were selected from anti-mouse CD8*α* antibody pretreated mice (cluster 2). Intravenous (i.v.) administration with vehicle or SNX281 was initiated on D0 as indicated below: Group 1: Vehicle (for SNX-281) + Isotype control antibody, 25 mg/kg; Group 2: Vehicle (for SNX281) + anti-CD8*α* antibody, 25 mg/kg; Group 3: SNX281 35mg/kg + Isotype control antibody, 25 mg/kg; Group 4: SNX281 35mg/kg + anti-CD8*α* antibody, 25 mg/kg; Group 5: SNX281 45mg/kg + Isotype control antibody, 25 mg/kg; Group 6: SNX281 45mg/kg + anti-CD8*α* antibody, 25 mg/kg, respectively. Vehicle and SNX281 were dosed via intravenous (i.v.) injection on D0 (single treatment) while antibodies were dosed via intraperitoneal injection on D1/5/10/15. Tumor volumes were measured three times per week for the duration of the study. 100*∼µ*L whole blood was collected for flow cytometry (FCM) analysis from mice in Groups 1, 2, 4 and 6 on Day 4. 100*∼µ*L anti-coagulant treated blood was collected for FCM analysis as described above. Red blood cells were removed using eBioscience 10xRBC lysis buffer (Cat : 00-4300-54) according to the manufacturer’s instructions.

Percent Relative Change of Body Weight (%RCBW) in mice treated with SNX281 was calculated according to the following formula:

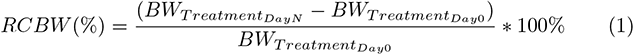

The percentage of circulating CD3+ CD8+ T-cells among CD45+ cells was measured by FCM 17 days after a single i.v dose.a). 100*µ*L anti-coagulant treated blood was collected for FCM analysis, 10 *µ*L MHC tetramer and other cell surface antibodies were added and incubated at 4°C for 40 min in the dark, red blood cells were removed using eBioscienceTM 10×RBC Lysis Buffer (Invitrogen), samples were re-suspended in 200 *µ*L Fixation/Permeabilization working solution (eBioscience, Cat : 00-5523-00) and incubated at 4*°*C for 40 min, cells were washed with 1xPermeabilization/Wash buffer and stained with Ki67-BV421 (Diluted in 1xPermeabilization/Wash buffer) at 4*°*C for 40 min. Stained samples were centrifuged and suspended in cell staining buffer, and analyzed on Thermo Fisher Attune NxT flow cytometer.

### 4.4 Chemistry

Tritiated cGAMP was synthesized via a bio-catalytic reaction in which recombinant cGAMP synthase preactivated with herring DNA was incubated with [3H]-ATP (Perkin Elmer) and [3H]-GTP (Perkin Elmer) overnight at 37*°*C. The reaction was then filtered to remove protein, and [3H]-cGAMP was purified by anion exchange chromatography.

Compounds 1-20 and 29-38 were prepared according to Schemes 1a-1c. The first step was a copper-catalyzed Ullmann-type coupling between the requisite aniline and the properly substituted 2-bromobenzoic acid or 2-iodobenzoic acid to form the corresponding diphenylamine. Representative reaction conditions include either Cu2O (0.05 eq), Cu (0.1 eq), and K2CO3 (1.2 eq), refluxed in DMF overnight. Variations in base, solvent, and copper source could be employed.

The second step was a cyclodehydration in H2SO4: H2O (10:1) at elevated temperature (80*°* C. or higher, overnight) to afford the substituted 10H-acridin-9-one. Other strongly acidic reagents, for example, polyphosphoric acid, were also effective for this purpose. In cases where the diphenylamine was synthesized from a starting aniline that did not possess an ortho-substituent, a mixture of regioisomers was obtained from the cyclodehydration. This mixture could be separated using normal phase silica gel chromatography.

Alkylation of the 10H-acridin-9-one was accomplished with ethyl bromoacetate and an excess of Cs2CO3 with heating in DMF (80*°* C., 2 hr). The ester of the bromoacetate, the base, and the solvent utilized in this step could be modified as needed. If the alkylation produced a putative mixture of N- and O-alkylated products, the ratio depended on the 10H-acridin-9-one substituents. These products were saponified as a mixture using metal hydroxide (NaOH, LiOH, or KOH) in a protic solvent, such as methanol or a THF/water mixture. Purification by extraction or reverse phase prep-HPLC delivered the desired substituted 10-carboxymethyl-9-acridinone.

### 4.5 Computational assays

#### 4.5.1 Protein model building, refinement, and ligand placement

Structures of the hSTING C-terminal domain were prepared for *in silico* applications from electron density by assigning bond orders, adding hydrogen atoms, and assigning appropriate protonation/tautomer states for the ligand and protein. Subsequently, hydrogen bond networks were optimized (pH set to 7.4) and a restrained energy minimization of only the hydrogen atoms was performed. To minimize overfitting during refinement with Phenix, a scan of the phenix.real space refine weight parameter was performed. To reduce local overfitting, ligand energy was monitored against the local map-model correlation as a function of the weight factor, and final weights were determined such that ligand energy values were no more than 0.1 log unit (1*∼*kcal/mol) greater than that observed for the lowest energy conformer. The same weights were tested for each structure, and the optimal weights were selected independently for each structure dataset collected.

#### 4.5.2 Grid Inhomogeneous Solvation Theory (GIST)

Molecular dynamics (MD) simulations of the C-terminal domain (CTD) of the human STING protein in explicit water were carried out with the program AMBER 18 Gõtz et al (2012). The GIST methodology calculates solvation entropy (Δ*S_solv_*) and solvation energy (Δ*E_solv_*), where both solute-water and water-water terms are readily computed. The spatial integrals in the inhomogeneous solvation theory expressions are approximated by discrete sums over voxels of a three-dimensional grid, in which the quantities on the grid are computed from MD simulation snapshots. More specifically, a spatial region G, which can include solute and solvent, is discretized into k voxels, having volumes *V_k_* at coordinates *r_k_*. The approximations inherent in this discretization scheme become exact in the limit *V_k_ >* 0, making smaller voxels more accurate given adequate sampling and convergence of solvent configurations. Details of the GIST calculations are described by Kurtzman and coworkers Ramsey et al (2016); **?**. Water thermodynamic data were processed using an in-house Python program and visualized using the open source version of PyMOL Schrödinger, LLC (2015).

#### 4.5.3 Functional Symmetry Adapted Perturbation Theory (F-SAPT)

Functional Group Symmetry-Adapted Perturbation Theory (F-SAPT) Parrish et al (2018) calculations were performed using the TeraChem Ufimtsev and Martinez (2009) quantum chemistry package run on graphic processing units (GPUs). F-SAPT calculations were run on individual frames from MD simulations of the CTD of the ligand bound hSTING protein in explicit water were carried out as described above. The functional group fragmentation patterns for the ligands were defined as variations on the 4-position of the acridone core and protein residues within 3*∼*^°^A of the ligand. Residual interaction energies were calculated as the sum of dispersion, exchange, and electrostatic terms computed for each subsampled frame of the MD trajectory. The average of ten frames is reported for each protein ligand pair.

#### 4.5.4 Virtual Compound Enumeration (STX-Novo)

STX-Novo is the in-house developed tool from Silicon Therapeutics used to perform reaction-based enumeration and generative modeling of novel small molecules. Input for STX-Novo is a SMILES string corresponding to a reference molecule, which serves as a starting point for the generation of new virtual molecules. Each new virtual molecule is the product of synthetic pathways linked to purchasable building blocks according to a list of discrete reaction templates. The reaction templates are defined as structural transformations representing a valid chemical reaction, encoded as a SMARTS string pattern. Given a list of reaction templates, *R*, and a list of purchasable building blocks, *B*, our goal is to enumerate valid synthetic pathways, *P*, which produces new molecules with a desired structure or function. This process of enforcing chemical reaction rules and the commercial availability of starting materials provides a practical approach to generative molecular modeling and improves the probability of successful synthetic outcomes for molecules evaluated *in silico*.

#### 4.5.5 ADMET Machine Learning (ML)

##### Graph Neural Networks

In the context of molecular machine learning, the featurization of molecular topology is paramount to effectively modeling the task at hand. For our ADMET machine learning models, molecules are modelled as undirected graphs, where each atom and bond can carry attributes reflecting their chemical nature from which complex chemical features can be learned. This can be formalized as a tuple of three sets,

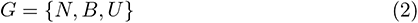

Here, *N* is the set of the nodes (atoms), *B* the set of bonds (edges), and *U* = *{u}* the universal (global) attribute.

A set of functions govern the discrete stages used in both training and inference of a graph neural network: initialization, propagation, and readout. Briefly, the most general description of the message-passing procedure during the propagation stage where node, edge, and global attributes *n, b, u* are updated according to:

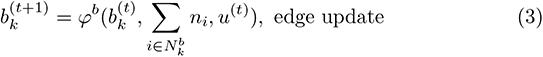

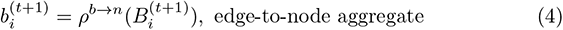

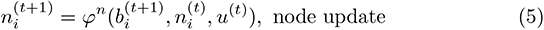

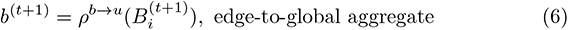

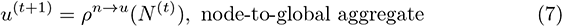

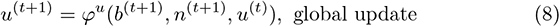

where 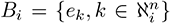 is the set of attributes of edges connected to a specific node, *B_i_* = *{e_k_, k ∈* 1, 2*, …, N^e^}* is the set of attributes of all edges, N is the set of attributes of all nodes.

##### Gaussian Process Regression

Gaussian processes (GP) are a Bayesian machine learning strategy that can learn nonlinear functions, can work with limited data, and enable principled incorporation of prior information. GPs enable a researcher to explicitly specify prior information encoding both a “baseline” prediction and corresponding uncertainty. A GP regressor is fully described by a mean function and a covariance function. For each compound-assay experiment, our mean function is specified as the upper limit of the assay format. The covariance function is set to a Gaussian, or squared exponential, kernel scaled by a constant prior k*_prior_* related to the prior uncertainty

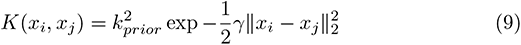

where ‖*·*‖_2_ denotes the *l*_2_-distance between feature vectors *x_i_* and *x_j_*. For the ADMET experiments, k*_prior_* is set to the upper limit of the assay. Each prediction takes the form of a scalar Gaussian distribution. We use the mean as the prediction value and the variance as the uncertainty estimate. We incorporate our graph neural network formalism into GPR by collapsing the adjendency matrix as input to a fixed dimensional kernel Wilson and Nickisch (2015). Therefore, we model the process that maps molecular graphs to physical properties, *f* : *G → y*, as a Gaussian process. We use the GP regressor implementation provided by the scikit-learn Pedregosa et al (2011) Python package.

#### 4.5.6 Pose Placement (Docking)

To prepare molecules generated by STX-Novo for 3D modeling and simulation, molecules were standardized and protonated at pH 7.4. Initial conformers were generated using default RDKit parameters using ETKDG **?**. Pose placement was performed using rDock v3.9 with core constraints defined as the maximum common substructure (MCS) between a new molecule and the template cocrystal structure Ruiz-Carmona et al (2014). Top scoring docking poses were selected for simulation preparation (unless the score was *>* 0, indicating no acceptable pose was discovered). Molecules were further parameterized and prepared for molecular dynamics simulations as previously described Lee et al (2020b).

#### 4.5.7 Molecular Dynamics (MD)

Molecular dynamics (MD) simulations of the C-terminal domain (CTD) of the human STING protein in explicit water were carried out with the program AMBER 18 Götz et al (2012). To evaluate pose stability and intermolecular interactions, three replicates of 100 nanosecond simulations were performed using the NPT ensemble (constant number of particles, pressure, and temperature). Ligand interactions/RMSD calculations were performed every 100 picoseconds.

#### 4.5.8 Relative Binding Free Energy Simulations on the Ligand Dimer (Dual-RBFE)

Relative binding free energy (RBFE) calculations were performed in AMBER as previously described Lee et al (2020b). In short, for each ligand pair, the alchemical transformation path was separated into 16 lambda windows and 5 ns of MD simulations were performed on each lambda window. However, the unique dimerization mechanism required a novel approach to binding free energy determination. We developed dual perturbation-RBFE (Dual-RBFE) to improve the overlap of phase space in our calculations by perturbing each copy of the ligand individually in a four leg thermodynamic cycle which allowed for better uncertainty quantification and hysteresis estimation, as shown in Figure 3. The workflow is thus defined as:

- Begin with the unperturbed copies of ligand *A* and *A^t^* bound (*AA^t^*) – Path 1: Perturb *A → B* with *A^t^* remaining unperturbed (*BA^t^*) – Path 2: Perturb *A^t^ → B^t^* with path 1 perturbed compound *B* bound (*BB^t^*)
- Only one simulation *A → B* is necessary for the solvent leg which is multiplied by two for the full thermodynamic cycle
- Cycle closure errors and calculation of ΔΔ*G_binding_* are computed based on the two paths

## Notes

### Competing Interest Statement

Authors are employees/shareholders of Silicon Therapeutics

### Summary of Updates

Figure revisions, grammar errors

## References

1. Ahlers LR, Goodman AG (2016) Nucleic Acid Sensing and Innate Immunity: Signaling Pathways Controlling Viral Pathogenesis and Autoimmunity. Current Clinical Microbiology Reports 3(3):132–141. https://doi.org/10.1007/s40588-016-0043-5, URL https://link.springer.com/article/10.1007/s40588-016-0043-5

2. Baguley BC, Ching LM (2002) DMXAA: An antivascular agent with multiple host responses. International Journal of Radiation Oncology Biology Physics 54(5):1503–1511. https://doi.org/10.1016/S0360-3016(02)03920-2

3. Beuming T, Che Y, Abel R, et al (2012) Thermodynamic analysis of water molecules at the surface of proteins and applications to binding site prediction and characterization. Proteins: Structure, Function, and Bioinformatics 80(3):871–883. https://doi.org/10.1002/prot.23244

4. Breiten B, Lockett MR, Sherman W, et al (2013) Water networks contribute to enthalpy/entropy compensation in protein-ligand binding. Journal of the American Chemical Society 135(41):15,579–15,584. https://doi.org/10.1021/JA4075776

5. Burdette DL, Vance RE (2013) STING and the innate immune response to nucleic acids in the cytosol. Nature Immunology 14(1):19–26. https://doi.org/10.1038/NI.2491

6. Chen L, Sun Y, Wang J, et al (2016) Differential regulation of the c-Myc/Lin28 axis discriminates subclasses of rearranged MLL leukemia. Oncotarget 7(18):25,208–25,223. https://doi.org/10.18632/ONCOTARGET.8199

7. Cournia Z, Allen B, Sherman W (2017) Relative Binding Free Energy Calculations in Drug Discovery: Recent Advances and Practical Considerations. Journal of Chemical Information and Modeling 57(12):2911–2937. https://doi.org/10.1021/acs.jcim.7b00564, URL https://pubs.acs.org/doi/10.1021/acs.jcim.7b00564

8. Diner EJE, Burdette DL, Wilson SC, et al (2013) No Title. Cell Reports 3(5):1355–1361

9. Eaglesham JB, Pan Y, Kupper TS, et al (2019) Viral and metazoan poxins are cGAMP-specific nucleases that restrict cGAS–STING signalling. Nature 566(7743):259–263. https://doi.org/10.1038/S41586-019-0928-6

10. Fuertes M (2011) Host type I IFN signals are required for antitumor CD8+ T cell responses through CD8alpha+ dendritic cells. J Exp Med 208(10):2005– 2016. https://doi.org/10.1084/jem.20101159

11. Fuertes MB, Woo SR, Burnett B, et al (2013) Type I interferon response and innate immune sensing of cancer. Trends in Immunology 34(2):67–73. https://doi.org/10.1016/J.IT.2012.10.004

12. Gao P, Ascano M, Wu Y, et al (2013a) Cyclic [G(2,5)pA(3,5)p] Is the Metazoan Second Messenger Produced by DNA-Activated Cyclic GMP-AMP Synthase. Cell 153(5):1094–1107. https://doi.org/10.1016/j.cell.2013.04.046, URL https://linkinghub.elsevier.com/retrieve/pii/S0092867413005229

13. Gao P, Ascano M, Zillinger T, et al (2013b) Structure-Function Analysis of STING Activation by c[G(2,5)pA(3,5)p] and Targeting by Antiviral DMXAA. Cell 154(4):748–762. https://doi.org/10.1016/j.cell.2013.07.023, URL https://linkinghub.elsevier.com/retrieve/pii/S0092867413008945

14. Gao P, Ascano M, Zillinger T, et al (2013c) XStructure-function analysis of STING activation by c[G(2,5) pA(3,5)p] and targeting by antiviral DMXAA. Cell 154(4). https://doi.org/10.1016/J.CELL.2013.07.023

15. Gkirtzimanaki K, Kabrani E, Nikoleri D, et al (2018) IFN*α* Impairs Autophagic Degradation of mtDNA Promoting Autoreactivity of SLE Monocytes in a STING-Dependent Fashion. Cell Reports 25(4):921–933. https://doi.org/10.1016/J.CELREP.2018.09.001

16. Gonugunta VK, Sakai T, Pokatayev V, et al (2017) Trafficking-Mediated STING Degradation Requires Sorting to Acidified Endolysosomes and Can Be Targeted to Enhance Anti-tumor Response. Cell Reports 21(11):3234–3242. https://doi.org/10.1016/J.CELREP.2017.11.061

17. Götz AW, Williamson MJ, Xu D, et al (2012) Routine microsecond molecular dynamics simulations with AMBER on GPUs. 1. generalized born. Journal of Chemical Theory and Computation 8(5):1542–1555. https://doi.org/10.1021/ct200909j, URL https://pubs.acs.org/doi/abs/10.1021/ct200909j

18. Hartenfeller M, Eberle M, Meier P, et al (2011) A collection of robust organic synthesis reactions for in silico molecule design. Journal of Chemical Information and Modeling 51(12):3093–3098. https://doi.org/10.1021/ci200379p

19. Hong Z, Mei J, Li C, et al (2021) STING inhibitors target the cyclic dinucleotide binding pocket. Proceedings of the National Academy of Sciences 118(24):e2105465,118. https://doi.org/10.1073/pnas.2105465118, URL http://www.pnas.org/lookup/doi/10.1073/pnas.2105465118

20. Hwang J, Kang T, Lee J, et al (2019) Design, synthesis, and biological evaluation of C7-functionalized DMXAA derivatives as potential human-STING agonists (7). URL www.rsc.org/obc

21. Jin L, Xu LG, Yang IV, et al (2011) Identification and characterization of a loss-of-function human MPYS variant. Genes and Immunity 12(4):263–269. https://doi.org/10.1038/GENE.2010.75

22. Kranzusch P (2015) Ancient origin of cGAS-STING reveals mechanism of universal 2,3 cGAMP signaling. Mol Cell 59(6):891–903. https://doi.org/10.1016/j.molcel.2015.07.022

23. Lee TS, Lin Z, Allen BK, et al (2020a) Improved Alchemical Free Energy Calculations with Optimized Smoothstep Softcore Potentials. Journal of Chemical Theory and Computation 16(9):5512–5525. https://doi.org/10.1021/acs.jctc.0c00237, URL https://pubs.acs.org/doi/10.1021/acs.jctc.0c00237

24. Lee TTS, Allen BB, Giese TT, et al (2020b) Alchemical Binding Free Energy Calculations in AMBER20: Advances and Best Practices for Drug Discovery. Journal of Chemical Information and Modeling 60(11):5595 – 5623. https://doi.org/10.1021/acs.jcim.0c00613

25. Martin GR, Henare K, Salazar C, et al (2019) Expression of a constitutively active human STING mutant in hematopoietic cells produces an Ifnar1-dependent vasculopathy in mice. Life Science Alliance 2(3). https://doi.org/10.26508/LSA.201800215

26. Meyer EA, Castellano RK, Diederich F (2003) Interactions with aromatic rings in chemical and biological recognition. Angewandte Chemie - International Edition 42(11):1210–1250. https://doi.org/10.1002/ANIE.200390319

27. Miao L, Qi J, Zhao Q, et al (2020) Targeting the STING pathway in tumor-associated macrophages regulates innate immune sensing of gastric cancer cells. Theranostics 10(2):498–515. https://doi.org/10.7150/THNO.37745

28. Novotná B, Vaneková L, Zavřel M, et al (2019) Enzymatic Preparation of 2-5,3-5-Cyclic Dinucleotides, Their Binding Properties to Stimulator of Interferon Genes Adaptor Protein, and Structure/Activity Correlations. Journal of Medicinal Chemistry 62(23):10,676–10,690. https://doi.org/10.1021/ACS.JMEDCHEM.9B01062

29. Pan BS, Perera SA, Piesvaux JA, et al (2020) An orally available non-nucleotide STING agonist with antitumor activity. Science 369(6506):eaba6098. https://doi.org/10.1126/science.aba6098, URL https://science.sciencemag.org/content/369/6506/eaba6098 https://science.sciencemag.org/content/369/6506/eaba6098.abstract https://www.sciencemag.org/lookup/doi/10.1126/science.aba6098

30. Parrish RM, Thompson KC, Martínez TJ (2018) Large-Scale Functional Group Symmetry-Adapted Perturbation Theory on Graphical Processing Units. https://doi.org/10.1021/acs.jctc.7b01053, URL https://pubs.acs.org/doi/abs/10.1021/acs.jctc.7b01053 https://pubs.acs.org/sharingguidelines

31. Pedregosa F, Varoquaux G, Gramfort A, et al (2011) Scikit-learn: Machine Learning in Python. Journal of Machine Learning Research 12(85):2825–2830. URL http://jmlr.org/papers/v12/pedregosa11a.html

32. Prabakaran T, Bodda C, Krapp C, et al (2018) Attenuation of c GAS - STING signaling is mediated by a p62/ SQSTM 1-dependent autophagy pathway activated by TBK1 . The EMBO Journal 37(8). https://doi.org/10.15252/EMBJ.201797858

33. Ramsey S, Nguyen C, Salomon-Ferrer R, et al (2016) Solvation thermodynamic mapping of molecular surfaces in ambertools: GIST. Journal of Computational Chemistry 37(21):2029–2037. https://doi.org/10.1002/jcc.24417

34. Rodríguez-García E, Olagüe C, Ríus-Rocabert S, et al (2018) TMEM173 Alternative Spliced Isoforms Modulate Viral Replication through the STING Pathway. ImmunoHorizons 2(11):363–376. https://doi.org/10.4049/IMMUNOHORIZONS.1800068

35. Rui Y, Su J, Shen S, et al (2021) Unique and complementary suppression of cGAS-STING and RNA sensing-triggered innate immune responses by SARS-CoV-2 proteins. Signal Transduction and Targeted Therapy 6(1). https://doi.org/10.1038/S41392-021-00515-5

36. Ruiz-Carmona S, Alvarez-Garcia D, Foloppe N, et al (2014) rDock: A Fast, Versatile and Open Source Program for Docking Ligands to Proteins and Nucleic Acids. PLOS Computational Biology 10(4):e1003,571. https://doi.org/10.1371/JOURNAL.PCBI.1003571, URL https://journals.plos.org/ploscompbiol/article?id=10.1371/journal.pcbi.1003571

37. Schrödinger, LLC (2015) The *{*PyMOL*}* Molecular Graphics System, Versioñ1.8

38. Shang G, Zhu D, Li N, et al (2012) Crystal structures of STING protein reveal basis for recognition of cyclic di-GMP. Nature Structural and Molecular Biology 19(7):725–727. https://doi.org/10.1038/NSMB.2332

39. Sintim HO, Mikek CG, Wang M, et al (2019) Interrupting cyclic dinucleotide-cGAS–STING axis with small molecules. MedChemComm 10(12):1999– 2023. https://doi.org/10.1039/C8MD00555A

40. Sivick KE, Desbien AL, Glickman LH, et al (2018) Magnitude of Therapeutic STING Activation Determines CD8+ T Cell-Mediated Anti-tumor Immunity 25(11):3074–3085. URL https://pubmed.ncbi.nlm.nih.gov/30540940/

41. Song L, Lee TS, Zhu C, et al (2019) Validation of AMBER/GAFF for Relative Free Energy Calculations (517):1–13. https://doi.org/10.26434/chemrxiv.7653434.v1, URL https://chemrxiv.org/articles/Validation_of_AMBER_GAFF_for_Relative_Free_Energy_Calculations/7653434/1

42. Sun L, Wu J, Du F, et al (2013) Cyclic GMP-AMP synthase is a cytosolic DNA sensor that activates the type I interferon pathway. Science 339(6121):786–791. https://doi.org/10.1126/SCIENCE.1232458

43. Ufimtsev IS, Martinez TJ (2009) Quantum chemistry on graphical processing units. 3. Analytical energy gradients, geometry optimization, and first principles molecular dynamics. Journal of Chemical Theory and Computation 5(10):2619–2628. https://doi.org/10.1021/CT9003004/SUPPL{_}FILE/CT9003004{_}SI{_}001.PDF, URL https://pubs.acs.org/doi/full/10.1021/ct9003004

44. Wang Y, Luo J, Alu A, et al (2020) CGAS-STING pathway in cancer biotherapy. Molecular Cancer 19(1). https://doi.org/10.1186/S12943-020-01247-W

45. Weiss JM, Guérin MV, Regnier F, et al (2017) The STING agonist DMXAA triggers a cooperation between T lymphocytes and myeloid cells that leads to tumor regression. OncoImmunology 6(10):e1346,765. https://doi.org/10.1080/2162402x.2017.1346765

46. Wilson AG, Nickisch H (2015) Kernel Interpolation for Scalable Structured Gaussian Processes (KISS-GP). 32nd International Conference on Machine Learning, ICML 2015 3:1775–1784. URL https://arxiv.org/abs/1503.01057v1

47. Woo S (2014) STING-dependent cytosolic DNA sensing mediates innate immune recognition of immunogenic tumors. Immunity 41(5):830–842. https://doi.org/10.1016/j.immuni.2014.10.017

48. Wu J, Zhao L, Hu H, et al (2020) Agonists and inhibitors of the STING pathway: Potential agents for immunotherapy. Medicinal Research Reviews 40(3):1117–1141. https://doi.org/10.1002/med.21649, URL https://onlinelibrary.wiley.com/doi/10.1002/med.21649

49. Yi G, Brendel VP, Shu C, et al (2013) Single Nucleotide Polymorphisms of Human STING Can Affect Innate Immune Response to Cyclic Dinucleotides. PLoS ONE 8(10):e77,846. https://doi.org/10.1371/journal.pone.0077846, URL http://www.ncbi.nlm.nih.gov/pubmed/24204993 http://www.pubmedcentral.nih.gov/articlerender.fcgi?artid=PMC3804601 https://dx.plos.org/10.1371/journal.pone.0077846

50. Zhang C, Shang G, Gui X, et al (2019) Structural basis of STING binding with and phosphorylation by TBK1. Nature 567(7748):394–398. https://doi.org/10.1038/s41586-019-1000-2, URL http://www.nature.com/articles/s41586-019-1000-2

51. Zhang X, Shi H, Wu J, et al (2013) Cyclic GMP-AMP Containing Mixed Phosphodiester Linkages Is An Endogenous High-Affinity Ligand for STING. Molecular Cell 51(2):226–235. https://doi.org/10.1016/j.molcel.2013.05.022, URL https://linkinghub.elsevier.com/retrieve/pii/S1097276513004097

52. Zhao Q, Manohar M, Wei Y, et al (2019) STING signalling protects against chronic pancreatitis by modulating Th17 response. Gut 68(10):1827–1837. https://doi.org/10.1136/GUTJNL-2018-317098

